# BAC Transgenic Expression of Human TREM2-R47H Remodels Amyloid Plaques but Unable to Reprogram Plaque-associated Microglial Reactivity in 5xFAD Mice

**DOI:** 10.1101/2023.08.03.551881

**Authors:** C.Y. Daniel Lee, Amberlene J. De La Rocha, Kellie Inouye, Peter Langfelder, Anthony Daggett, Xiaofeng Gu, Lu-Lin Jiang, Zoe Pamonag, Raymond G. Vaca, Jeffrey Richman, Riki Kawaguchi, Fuying Gao, Huaxi Xu, X. William Yang

## Abstract

**Background:** Genetic study of late-onset Alzheimer’s disease (AD) reveals that a rare Arginine-to-Histamine mutation at amino acid residue 47 (R47H) in Triggering Receptor Expressed on Myeloid Cells 2 (TREM2) results in increased disease risk. TREM2 plays critical roles in regulating microglial response to amyloid plaques in AD, leading to their clustering and activation surrounding the plaques. We previously showed that increasing human *TREM2* gene dosage exerts neuroprotective effects against AD-related deficits in amyloid depositing mouse models of AD. However, the *in vivo* effects of the R47H mutation on human TREM2-mediated microglial reprogramming and neuroprotection remains poorly understood.

**Method:** Here we created a BAC transgenic mouse model expressing human TREM2 with the R47H mutation in its cognate genomic context (BAC-TREM2-R47H). Importantly, the BAC used in this study was engineered to delete critical exons of other TREM-like genes on the BAC to prevent confounding effects of overexpressing multiple TREM-like genes. We crossed BAC-TREM2- R47H mice with 5xFAD [1], an amyloid depositing mouse model of AD, to evaluate amyloid pathologies and microglial phenotypes, transcriptomics and *in situ* expression of key *TREM2*-dosage dependent genes. We also compared the key findings in 5xFAD/BAC-TREM2-R47H to those observed in 5xFAD/BAC-TREM2 mice.

**Result:** Both BAC-TREM2 and BAC-TREM2-R47H showed proper expression of three splicing isoforms of *TREM2* that are normally found in human. In 5xFAD background, elevated TREM2-R47H gene dosages significantly reduced the plaque burden, especially the filamentous type. The results were consistent with enhanced phagocytosis and altered NLRP3 inflammasome activation in BAC- TREM2-R47H microglia in vitro. However, unlike TREM2 overexpression, elevated TREM2- R47H in 5xFAD failed to ameliorate cognitive and transcriptomic deficits. *In situ* analysis of key *TREM2*-dosage dependent genes and microglial morphology uncovered that TREM2-R47H showed a loss-of-function phenotype in reprogramming of plaque-associated microglial reactivity and gene expression in 5xFAD.

**Conclusion:** Our study demonstrated that the AD-risk variant has a previously unknown, mixture of partial and full loss of TREM2 functions in modulating microglial response in AD mouse brains. Together, our new BAC-TREM2-R47H model and prior BAC-TREM2 mice are invaluable resource to facilitate the therapeutic discovery that target human TREM2 and its R47H variant to ameliorate AD and other neurodegenerative disorders.

## BACKGROUND

Genome-wide association studies (GWAS) of late-onset Alzheimer’s disease (AD) have revealed more than 20 genome-wide significant disease-risk associated loci [2]. Interestingly, a large number of AD GWAS locus genes are related to innate immunity and are selectively expressed in microglia in the brain and myeloid cells in the peripheral tissues [3–7]. Thus, human genetics highlights microglia and the function of several key genes in this cell type in AD pathogenesis [6, 7].

Among the AD GWAS genes, variants of Triggering Receptor Expressed in Myeloid Cell 2 (TREM2), e.g. TREM2-R47H, are particularly important as they confer 2-4 fold increase in AD risk, and such magnitude of increase is second only to APOE4 [2, 8, 9]. TREM2 is a type I membrane protein with an extracellular receptor domain and a short intracellular signaling domain, and it is expressed exclusively in microglia in the brain [7, 10]. TREM2 signals through a binding partner DAP12 (encoded by the gene *TYROBP*) and activates downstream signaling pathways through kinases including SYK and PI3K [11]. TREM2-DAP12 signaling elicits diverse effects in microglia, including enhancing phagocytosis [12], suppressing proinflammatory Toll-like receptor signaling [13, 14], promoting microglia survival [15, 16], and activating disease-associated microglial (DAM) gene expression signatures in neurodegenerative and neuroinflammatory diseases [17, 18].

Loss-of-function (LOF) variants in TREM2 or DAP12 (encoded by *TYROBP* gene) results in polycystic lipomembranous osteodysplasia with sclerosing leukoencephalopathy (PLOSL), also known as Nasu-Hakola disease (NHD), which is characterized by recurrent bone cysts, progressive demyelination and axonal degeneration in the brain, and presenile dementia [19, 20]. In mouse models of familial AD, *Trem2*-deficiency impairs microglial clustering at the amyloid plaques, dampens microglial morphological and transcriptomic activation (e.g. DAM signatures), and enhances fibrillary Aβ plaques and dystrophic neurites [15, 21]. In contrast to the *Trem2* deficiency, we previously showed that elevated human *TREM2* gene dosage (but not any other TREM-like genes) using Bacterial Artificial Chromosome (BAC)-transgenic mouse models (BAC-TREM2) ameliorates amyloid burden, remodels plaques from filamentous to inert types, reprograms plaque-associated microglial morphology and transcriptomic activation signatures, and rescues dystrophic neurites and cognitive deficits in mouse models of AD [22]. These findings suggested that increase the expression of TREM2 in a human genomic context alters microglial responsivity and confers multiple benefits in ameliorating disease in amyloid AD mouse models.

The rare homozygous *TREM2-R47H* cases have been reported to present classic AD phenotypes with a relatively early age of onset yet without bone cysts, suggesting that the R47H mutation is not a full LOF mutation compared to the *TREM2* mutations in NHD [23]. The R47H mutation is located in the extracellular domain of TREM2 [24] and has been shown to interfere with its ligand binding capability to zwitterionic lipids [15, 25–27] and APOE [26–28]. In AD patient brains, the *TREM2-R47H* allele was shown to result in fewer microglia surrounding the plaque, an increase in filamentous amyloid plaques, and worsening of dystrophic neurites and Tau pathology [29]. Several mouse models have been developed to study the effects of R47H knockin variants in murine *Trem2* [30, 31] or transgenically-expressed human *TREM2* [32, 33]. The initial *Trem2-R47H* knockin mouse model exhibited loss-of-function phenotype but was attributed to the reduction of full-length *Trem2* mRNA due to cryptic splicing introduced by the CRISPR engineering of the allele [30, 31]. A new *Trem2-R47H* knockin allele without the cryptic splicing transcripts was recently generated, which showed proper Trem2-dependent inflammatory response with treatment of cuprizone, a demyelinating agent [34]. Crossing this *Trem2-R47H* “normal splice site” allele to 5xFAD mice showed an early *Trem2* loss-of-function phenotype including reduced plaque-associated microglia and increased dystrophic neurites at 4 months of age (4m), phenotypes similar to *Trem2* null mutants in such AD model [15]. However, these changes resolved at 12 months of age (12m) and 5xFAD with Trem2-R47H showed an interferon signature and upregulation of ribosomal gene expression [34]. Humanized TREM2-R47H mice, expressed from a *TREM2* BAC transgene in a murine *Trem2* null background, also showed a lack of plaque-associated microglia clustering and activation suggesting a loss-of-function phenotype, which are consistent with LOF phenotype [33]. However, this transgene was not engineered to delete the other TREM-like genes on the BAC [33]. Therefore, any effects due to the co-expression of other human TREM-like genes from the same BAC transgene cannot be fully discerned. Recently, wildtype or AD-associated *TREM2-R47H* cDNA were used to replace the endogenous murine *Trem2*, and the *TREM2-R47H* but not control TREM2 knockin allele were shown to enhance inflammation in a Tauopathy mouse model through activation of Akt signaling in microglia, suggesting a possible gain-of-function phenotype [32]. However, the *TREM2-R47H* knockin allele appear to show multiple beneficial effects in a Tau transgenic model [35]. Thus far, the impact of *Trem2-R47H (or TREM2-R47H)* knockin or human TREM2-R47H in AD mouse models consistently show an overall LOF phenotypes in the amyloid AD models but their effects in Tauopathy models appear to be mixed. However, these prior studies could not delineate the granularity of differential microglia-plaque responses modulated by TREM2-R47H compared to the wildtype TREM2, especially in a human TREM2 DNA/RNA/protein context.

In the current study, we generated BAC-TREM2-R47H mice to increase the gene dosage of human *TREM2-R47H* variant and studied its effects in modifying microglial and amyloid phenotypes in 5xFAD mice. We compared overexpressing BAC-TREM2-R47H and BAC-TREM2 in the 5xFAD background to draw conclusions on the differential impacts of R47H versus common variants of human TREM2 on a range of AD-related phenotypes *in vivo*. Our study showed TREM2-R47H partially retained wildtype TREM2 function in reducing and remodeling the amyloid plaque pathology and enhancing microglial phagocytosis *in vitro*. However, the mutant TREM2-R47H failed to retain the TREM2 functions in reprogramming morphology and gene activation of microglia surrounding the amyloid plaques. Thus, our studies suggest that human *TREM2-R47H* variant loses some, but not all, of its functions in the microglia that are crucial to modify disease-related phenotypes in an amyloid AD mouse model; and also provides an *in vivo* human genomic BAC-TREM2-R47G model system to study pathogenesis and therapeutics of AD.

## METHODS

### Generation of BAC Transgenic Mice

To introduce the R47H mutation, we re-engineered the TREM2 BAC used in our previous study [22] by two sequential modification steps using RecA-based shuttle vector plasmids described previously [36–38]. We first replaced the exon 2 of TREM2, which contains the codon (CGC) for the R47 residue, with a spacer sequence. In the second step, this spacer sequence was replaced by the exon 2 with the codon (CGC) for R47 substituted by CAC, which encodes histamine, producing an identical BAC transgene construct to BAC-TREM2 except for this single SNP. The modified TREM2-R47H BAC DNA was prepared according to our published protocols and microinjected into FvB fertilized oocytes. BAC-TREM2-R47H mice were maintained in the FvB/NJ background.

### Animal breeding and husbandry

5xFAD mice (C56BL/6J background) were purchased from the Jackson Laboratory (MMRRC) and crossed to BAC-TREM2 mice (FvB/NJ background). Thus, all genotypes of mice used in the current study were generated and analyzed in the F1 hybrid background (C56BL/6J;FvB/NJ F1), which is suitable for phenotypic study of genetically engineered mutant mice (Silva et al., 1997). Animals were housed in standard mouse cages under conventional laboratory conditions, with constant temperature and humidity, 12h/12h light/dark cycle and food and water ad libitum. All animal studies were carried out in strict accordance with National Institutes of Health guidelines and approved by the UCLA Institutional Animal Care and Use Committees.

### Tissue collection and sample preparation

Mice were anesthetized with pentobarbital and perfused with DEPC-PBS. Brains were bisected. The right hemispheres were immediately submerged in ice-cold DEPC-PBS for dissecting out cortices, hippocampi and other brain regions. Dissected tissues were snap frozen in dry ice and stored in -80°C before further processing. The left hemispheres were fixed in 4% PFA/DEPC-PBS overnight followed by submergence in 30% sucrose before freezing. Coronal sections (40 µm) were obtained using a Leica cryostat and stored in cryopreserve solution (250mM sucrose, 3.3 mM MgCl2 in 1:1 (vol/vol) glycerol/0.1M PBS) at -20°C.

For preparing the samples for RNA sequencing and biochemistry, dissected brain tissues were homogenized and aliquoted as described previously [39, 40]. In brief, cortical and hippocampal tissues from one hemisphere were homogenized in tissue homogenization buffer (250 mM sucrose, 20 mM Tris at pH 7.4, 1 mM EDTA, and 1 mM EGTA in DEPC-treated water) and centrifuged at 5000 x g for 10 min at 4°C. Supernatants were aliquoted and stored at -80°C.

### Immunohistochemistry and image analysis

Coronal sections were blocked in the blocking buffer (3% BSA, 2% normal goat serum and 0.3% Triton X-100 in PBS) for 1 hour at room temperature and then incubated with primary antibodies at 4°C overnight. Incubation in secondary antibodies was performed for 2h at room temperature before mounting on slides with Prolong Diamond anti-fade mountant (ThermoFisher). Aβ plaques were visualized by ThioS staining or by immunostaining using anti-Aβ antibodies 6E10. For plaque number and categorization, 3 matched coronal sections/mouse spacing out across 1 mm (0.5 mm apart) were stained with 6E10 and ThioS. Z-stack 20x images covering the whole associate and visual cortex on the sections were taken on an Andor DragonFly confocal microscope and analyzed using ImageJ. Pixels with < 1% of max intensity were discarded as background and were not counted as a part of the plaque. All images were preprocessed using the same threshold setting prior to analysis. For microglial morphology, Z-stack 60x images of the plaque-associated microglia were collected from matched cortical regions. A total of 190 plaques with 1,305 plaque-associated microglia from four gender matched animals were analyzed using the FilamentTracer feature in Imaris (Bitplane). The image acquisition and quantification were performed in a blinded manner.

### RNA purification and mRNA sequencing

Total RNA was extracted using RNeasy kit (Qiagen). Library preparation and RNA sequencing were performed by the UCLA Neuroscience Genomics Core (UNGC). Libraries were prepared using the Illumina TruSeq RNA Library Prep Kit v2 and sequenced on an Illumina HiSeq4000 sequencer using strand-specific, paired-end, 69-mer sequencing protocol to a minimum read depth of 30 million reads per sample. Reads were aligned to mouse genome mm10 using the STAR aligner [41] with default settings. Human-specific *TREM2-R47H* reads were obtained by aligning to the human reference genome (build GRCh38) reads that failed to align to the mouse genome (build mm10). Mouse-specific *Trem2* reads were obtained in similar way. Mapped reads were quantified by HTSeq [42]. Read counts were divided by the library size per million to determine the counts per million (CPM). Homer [43] makeTagDirectory (parameters: - format sam -flip -sspe) and makeUCSCfile (parameters: -fragLength given -o auto -raw) functions, bedtools [44] and bedGraphToBigWig tools were used to create CPM bigwig tracks for visualization onto the UCSC genome browser.

### Differential expression analysis

For outlier removal, we retained mRNA profiles whose CPM is 1 or more in at least one-quarter of the samples and transformed the raw counts using variance stabilization. Outlier samples were removed as described [22], using the Euclidean distance-based sample connectivity Z.k threshold of -5. This procedure resulted in the removal of a single WT sample.

For DE testing and network analysis, we used individual observation weights constructed as follows. Tukey bi-square-like weights λ [45] are calculated for each (variance-stabilized) observation *x*, as

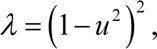

where *u* = min(1, *x* − *m* / (9*MAD*)), and *m* and *MAD* are median and median absolute deviation of the observations of the gene.

For each gene, *MAD* is adjusted such that (1) 10^th^ percentile of the weights *λ* is at least 0.1 (that is, the proportion of observations with coefficients <0.1 is less than 10%) [46] and (2) for each individual genotype, 40^th^ percentile of the weights *λ* is at least 0.9 (that is, at least 40% of the observation have a high coefficient of at least 0.9).

DE testing was carried out in R using package DESeq2 [22] version 1.28.1.

DESeq2 models observed counts using Negative Binomial Generalized Linear Models with dispersion estimated from data. Wald test was used to for significance calculations, and independent filtering was disabled. For differential expression testing between genotypes, sex was used as a covariate. Genotype-sex interactions were tested using models with genotype × sex terms (with genotype and sex turned into binary indicator variables).

For each genotype contrast or interaction, DE tests result in gene-wise Z statistics (fold changes divided by their standard errors). We assess the overall similarity of genome-wide effects of two genotype contrasts using a scatter plot of their gene-wise DE Z statistics. A positive linear trend (correlation) indicates that the effects of the two genotype contrasts are broadly similar, whereas a negative correlation indicates broadly opposing effects. Although other measures are possible, the correlation value can be used as a measure of similarity. We use the genome-wide correlations of DE Z statistics for 5xFAD vs. WT and 5xFAD vs. 5xFAD/R47H to define a measure of genome-wide rescue. The we refer to the corresponding scatterplot as the “rescue/exacerbation” plot.

We define significantly rescued and exacerbated genes as those whose DE for 5xFAD vs. WT is significant (FDR<0.1), DE for 5xFAD/R47H vs. 5xFAD is significant (FDR<0.1) and the direction of DE in these two sets is opposite (rescue) or same (exacerbation).

### Splicing analysis of TREM2

Reads were classified to five categories (human, mouse, both, ambiguous and neither) by Xenome (ver1.0.0). Reads classified as human reads were aligned to human genome (Hg38) by STAR aligner (ver2.6) using two pass alignment methods. Alignment data are sorted and indexed by samtools (ver1.7). rMATS (ver4.0.2) was used to detect alternative splicing events. Splicing variants were visualized by Sashimi plots using rmats2sashimiplot (v2.0.0).

### Real-time RT-PCR

For the qPCR analyses, 1 μg of total RNA was reversed transcribed using the QuantiTect Reverse Transcription kit (ThermoFisher, #4387406). Fast SYBR Green Master Mix (Applied Biosystems) was used to perform real-time PCR on a QuantStudio 5 thermocycler (Applied Biosystems). The abundance of each transcript was normalized to GAPDH as ΔΔCt. Doubling efficiency was also calculated.

### RNAscope HiPlex fluorescent in situ hybridization assay and image analysis

Multiplex in situ hybridization was performed on coronal sections (40 μm) using RNAscope HiPlex according to the manufacturer’s protocol for fresh-frozen tissue (ACD/Biotechne). The HiPlex together with immunofluorescent multiplex assay allows multiplex detection for up to 15 targets (including mRNA and protein targets) on a single tissue section. Briefly, sections were fixed in 4% paraformaldehyde, dehydrated with 50%, 70%, 100% ethanol, then treated with protease. In HiPlex, the probes for human TREM2 (Cat No. 420498-T3) and mouse Tyrobp (Cat No. 408191-T2), Cst7 (Cat No. 498711-T7), Spp1 (Cat No. 435191-T6), Gpmnb (Cat No. 489511-T9) and Atp6v0d2 (Cat No. 818041-T11) were used. The sections were incubated with pooled HiPlex probes all together. The first group of genes (Tyrobp, Spp1 and Gpnmb) were visualized by a series of amplification and detection solutions. Cell nuclei were counterstained with Hoechst 33342 (Sigma) and samples were mounted. Imaging was performed using an Andor DragonFly spinning-disk confocal microscope with a 30x objective. After imaging, the coverslips were removed by submerging the slides in PBS. Sections were treated with TCEP solution to cleave fluorophores, and then moved on to the next round (for hTREM2, Cst7 and Atp6v0d2) of the amplification/detection/imaging procedures. After RNAscope imaging, samples were proceeded to immunostaining to visualize microglia (Iba1+) and plaques (using ThioS and 6E10). Images from all rounds of staining were aligned and analyzed for the gene expression in plaque-associated microglia using Imaris.

### Primary Microglial Culture and phagocytosis assay

Primary microglia were isolated from the brains of neonatal mice at postnatal days 2-3 using a mild trypsinization protocol as previously described [47, 48]. For phagocytosis assay, purified microglia were maintained in DMEM/F-12 (ThermoFisher) containing 2% heat-inactivated fetal bovine serum (FBS) and 1% penicillin/streptomycin plated at a density of 250,000 cells/well in 24-well plates for 3-5 days before further experiments. The culture media was replenished with serum-free DMEM/F12 overnight. Cells were incubated with BSA (0.5 mg/ml in PBS)-preblocked microsphere (1 µm, Alexa 488-conjugated; ThermoFisher) for 30 min, followed by extensive washing with PBS and fixation with 4% paraformaldedyde. After fixation, cells were washed with PBS and collected for analysis using an LSR II flow cytometer (BD Biosciences).

### oAβ preparation

Aβ oligomers (oAβ) were prepared as described previously (Zhao et al, 2018). Synthetic human Aβ1-42 peptides (Anaspec) were dissolved in hydroxyfluroisopropanol (HFIP) and subsequently dried using a SpeedVac system. The lyophilized Abeta was dissolved in dimethyl sulfoxide (DMSO) at concentration of 2.2 mM, sonicated for 10 min and diluted in PBS to 100 μM. oAβ was generated by incubating the Aβ/PBS solution at room temperature for 48 h.

### Inflammasome activation and quantification

Isolated microglia were seeded on poly-D-lysine-coated coverslips at the density of 50,000 – 100,000 cells/well in the 24-well plates for overnight and treated with oAβ at a final concentration of 5 μM for 24 h. After the oAβ treatment, cells were fixed in 4% PFA at RT for 15 min and blocked in 5% BSA (dissolved in 0.5% Triton X-100-containing PBS) for 30min. The cells were then incubated with the following primary antibodies: NLRP3 (1:100, Adipogen, cat# AG-20B-0014-C100) and ASC (1:100, Cell Signaling Technologies, cat# 67824) at 4°C for overnight. The fluorophore-conjugated secondary antibodies were applied at RT for 1 hr: donkey anti-mouse IgG-Alexa Fluor 647 (Thermo, cat# A-31571, 1:1000), donkey anti-rabbit IgG-Alexa Fluor 555 (Thermo, cat# A-31572, 1:1000) and DAPI. After washing in PBS, the cover glass was mounted with Fluoromount-G mounting medium (Thermo, cat# 00-4958-02). The fluorescent images were taken using a Zeiss confocal microscope (LSM 710) and analyzed by ImageJ.

### Behavioral tests

Contextual fear conditioning test was performed on 7 months old mice as describe previously [22] and conducted in the Behavioral Testing Core (BTC) at UCLA. In brief, mice were handled daily for a week prior to the behavior test. In the training phase, mice were placed individually in the conditioning chamber to explore the environment freely for 2 min before the first unconditioned stimulus (US: 0.75 mA, 2 s) was delivered. The animals were exposed to 2 US’s with an intertrial interval of 3 min. After the last shock, the mice were left in the chamber for another 1 min and then placed back in their home cages. Retention tests were performed 24 hours later. Each mouse was returned to the same chamber for measuring the percent of time frozen and number of freezes. No shocks were given during the test session. Both training and testing procedures were videotaped, and the freezing behavior was measured by an automated tracking system (Med Associates).

### Statistical analyses

Statistics for transcriptomic and RNAscope analyses were described as above. Other quantitative results, unless otherwise specified, were analyzed using one-way ANOVA with Turkey post-hoc analysis to determine the *p* value. Morphological and behavioral studies were subjected to Power analysis to determine the biological replicates (*n*) needed to reach >80% confidence level.

### Data and software availability

RNA-seq data has been deposited within the Gene Expression Omnibus (GEO) repository (www.ncbi.nlm.nih.gov/geo). The accession number is pending updated.

## RESULTS

### Generation of BAC-TREM2-R47H mice to express the AD-risk allele of TREM2 gene under endogenous human genomic regulatory elements

To study the effect of the R47H mutation of TREM2 on AD pathogenesis, we used a previously engineered human TREM2 BAC with deletion of key exons in three other TREM2-like genes (i.e. *TREML1*, *TREML2* and *TREML4*) on the BAC [22]. We performed homology recombination-based BAC modification to introduce a G-to-A single nucleotide mutation, which encodes an R-to-H mutation in the residue 47 of TREM2 (Fig. 1a; [36]). The resulting BAC was used for pronuclear injections in the FvB/NJ embryos to generate BAC-TREM2-R47H transgenic founders. Three independent founders (A, C and D) gave germline transmission of their transgenes. We selected the BAC-TREM2-R47H line A (referred to as BAC-TREM2-R47H hereafter) and used genomic qPCR to estimate the transgene copy number to be one copy, which is the same as BAC-TREM2 mice [22]. Baseline expression of human *TREM2-R47H* mRNA in BAC-TREM2- R47H mice was also comparable to that of *TREM2* in BAC-TREM2 mice (Fig. 1b). The transcription of endogenous murine *Trem2* was not altered by the expression of human *TREM2- R47H* (Fig. 1c).

**Figure 1.**
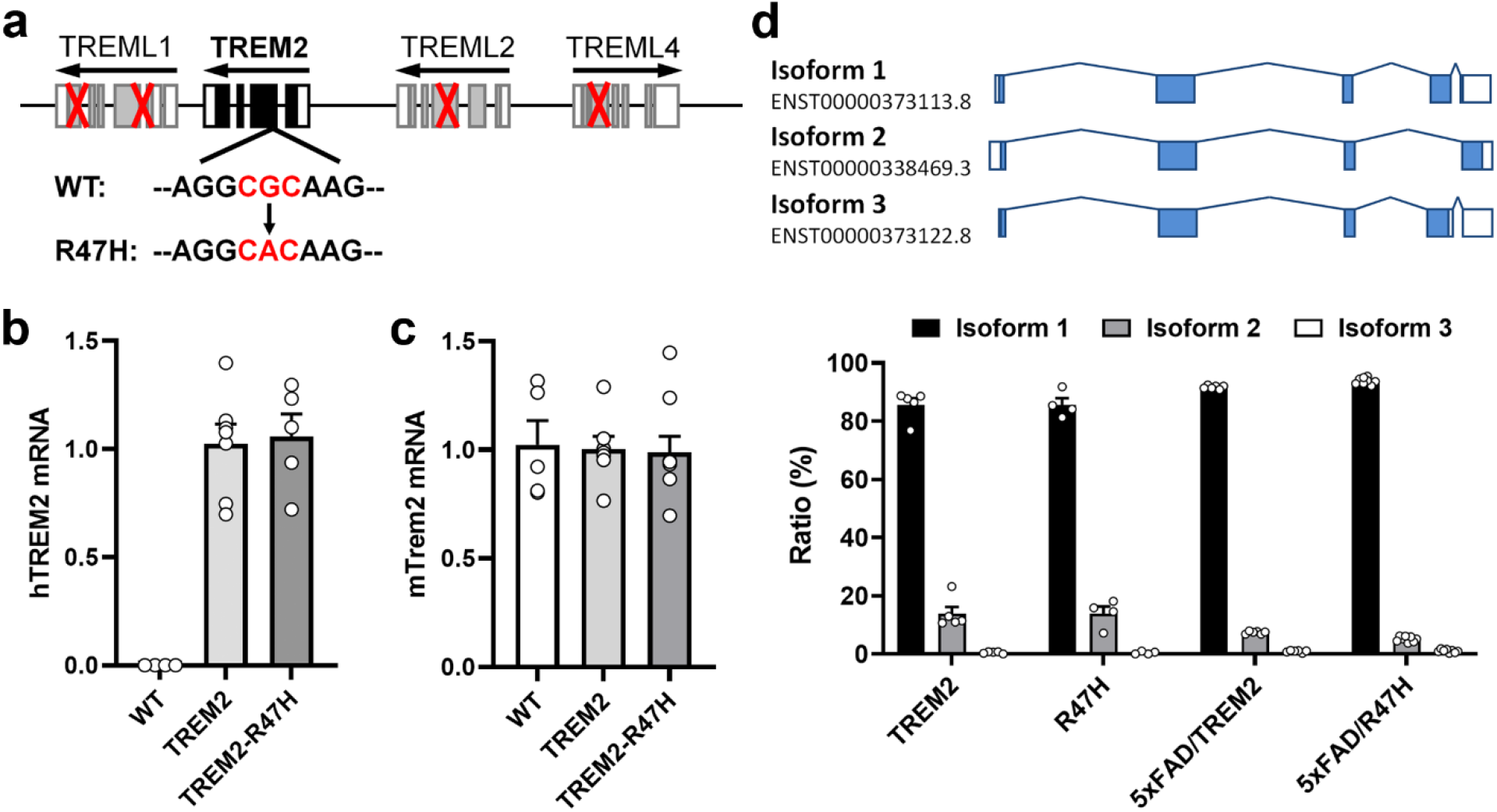
Generation and characterization of BAC-TREM2-R47H mice. (**a**) Schematic representation of the genetic modification of the TREM2 BAC to introduce the R47H mutation. Red crosses indicate the deleted exons in TREM-like genes in the BAC construct. (**b-c**) Real-time PCR was performed to quantify the levels of the human TREM2 (**b**) and mouse Trem2 transcripts (**c**) in the cortex of 2 months old mice. Genotypes of the mice (WT, BAC- TREM2 and BAC-TREM2-R47H) are indicated under the bars. (**d**) The splicing diagram of the three TREM2 isoforms are depicted in the top panel. TPM (Transcripts Per Kilobase Million) of the transcripts specifically assigned to each isoform was used to calculate the proportion of each isoform and plotted in the bar graph. *n* = 5-6.

Previous studies showed certain *Trem2-R47H* knockin alleles [30, 31], but not all [34], showed cryptic splicing resulting in nonsense mediated decay (NMD) of its mRNA transcripts. We assessed TREM2 alternative splicing patterns in BAC-TREM2 and BAC-TREM2-R47H mice and their crosses to 5xFAD for the presence of three human *TREM2* splice isoforms found in AD and control brains [49]. Isoform 1 (ENST00000373113) is the canonical form, encoding full-length TREM2 that includes extracellular, transmembrane, and intracellular domains. Isoform 2 (ENST00000338469) lacks exon 4 containing the transmembrane domain and therefore encodes a soluble and secreted *TREM2* isoform (sTREM2). Isoform 3 (ENST00000373122) is alternatively spliced exon 4, resulting in a premature Stop codon and a shortened transmembrane domain. By mapping cortical RNA-seq reads to the human genome assembly (see Methods), we found all three correctly spliced human *TREM2* isoforms in both BAC-TREM2 and BAC-TREM2-R47H in the WT and 5xFAD backgrounds (Fig. 1d; Fig. S1). Isoform 1 was the predominant isoform expressed in mice carrying BAC-TREM2 and BAC-TREM2-R47H transgenes, and isoform 2 (a source of sTREM2) and isoform 3 were the 2^nd^ and 3^rd^ most abundant splicing isoforms (Fig. 1d). In summary, BAC-TREM2-R47H and BAC-TREM2 transgenes confer the proper expression of all three human TREM2 splicing isoforms in the cortices of 5xFAD and control mice.

### Increased TREM2-R47H gene-dosage reduces amyloid plaque load and remodels amyloid pathology types

We next tested whether BAC-TREM2-R47H could exert similar plaque modifying effects as the wildtype BAC-TREM2 in 5xFAD [22]. We found that overexpression of TREM2-R47H significantly reduced Thioflavin S (ThioS)-positive amyloid plaques compared with 5xFAD at 7 months (7m) old (Fig. 2a, 2b, 2d). However, the magnitude of plaque load reduction is significantly smaller than that in 5xFAD/BAC-TREM2 mice (Fig. 2c, 2d). We next measured the levels of Aβ42 species in cortical samples of 5xFAD/BAC-TREM2-R47H and 5xFAD littermates. We found a significant reduction of the soluble Aβ42, but not insoluble Aβ42, in 5xFAD/BAC-TREM2-R47H cortical lysates compared to those from 5xFAD (Fig. 2e, 2f). These results suggest that TREM2- R47H partially retains such protective function compared to TREM2 [22]. Similar to our previous finding that elevated *TREM2* gene dosage remodeled the plaques and reduced the neurotoxic filamentous form of amyloid plaques [22], we also found a significant decrease of filamentous plaques in 5xFAD/BAC-TREM2-R47H mice compared to 5xFAD (Fig. 2g). However, only the 5xFAD mice overexpressing wildtype TREM2, but not TREM2-R47H, also showed a significant decrease in compact plaques and increase in inert plaques (Fig. 2g). Together, we concluded that *TREM2-R47H* expressed from a human BAC transgene partially retains the effects of wildtype *TREM2* gene-dosage-dependent effects in reducing amyloid plaque load and remodeling amyloid plaque types.

**Figure 2.**
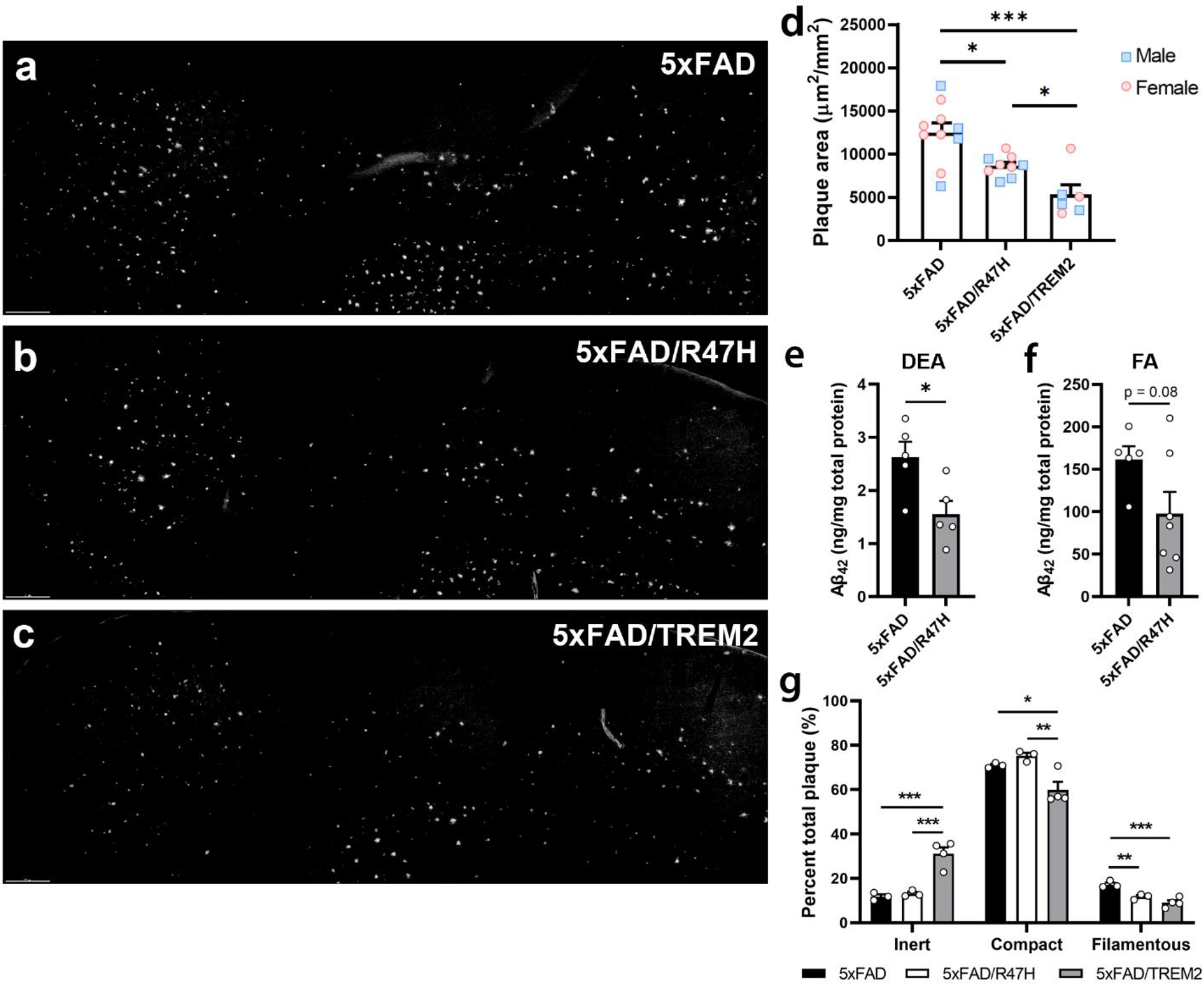
Overexpressing TREM2-R47H reduced amyloid burden and pathology in 5xFAD mice. (**a-d**) Representative images of matched brain sections from 7-month-old 5xFAD (**a**), 5xFAD/R47H (**b**) and 5xFAD/TREM2 mice (**c**) were stained with ThioS to visualize the amyloid plaques in the cortex. Z-stack confocal images (30X) were utilized to measure total plaque area in the field using ImageJ. The results are presented as ThioS^+^ plaque area (μm^2^) per mm^2^ of the cortical area (**d**). *n* = 5-10 per genotype. Bar = 200μm. (**e-f**) The soluble and insoluble fractions (i.e. DEA and FA fractions, respectively) for Aβ in the cortical samples were measured by ELISA. *n* = 5 per genotype. (**g**) Brain sections from 7-month-old male 5xFAD, 5xFAD/R47H and 5xFAD/TREM2 mice were stained with ThioS and an anti-Aβ antibody (6E10). Z-stack confocal images (30X) were utilized to quantify 3 different types of plaques using ImageJ (J). A total of 1,028 plaques were analyzed. Data are represented as mean ± SEM. *n* = 4 per genotype, \**p*<0.05, \*\**p* < 0.01, \*\*\**p* < 0.001 compared to 5xFAD or otherwise indicated.

### Increased TREM2-R47H gene-dosage enhances baseline phagocytic activity and attenuate oligomeric Aβ-induced inflammasome activation in primary microglia

TREM2 promotes microglial phagocytosis of several substrates, including apoptotic neurons [12], Aβ [50] and myelin debris [51]. We previously reported that overexpressing *TREM2* enhanced microglial phagocytic capacity, while *Trem2* deficiency significantly impaired it [22]. We next evaluated primary microglia from neonatal BAC-TREM2, BAC-TREM2-R47H or WT mice for their general phagocytic capacity (i.e. engulfment of polystyrene microbeads), and found that a significant enhancement of phagocytic activity in both BAC-TREM2 and BAC-TREM2- R47H microglia compared to WT microglia (Fig. 3a). No significant difference was observed in this readout for microglia expressing BAC-TREM2 or BAC-TREM2-R47H, suggesting TREM2- R47H retaining the TREM2 function in boosting microglial general phagocytic capacity *in vitro*.

**Figure 3.**
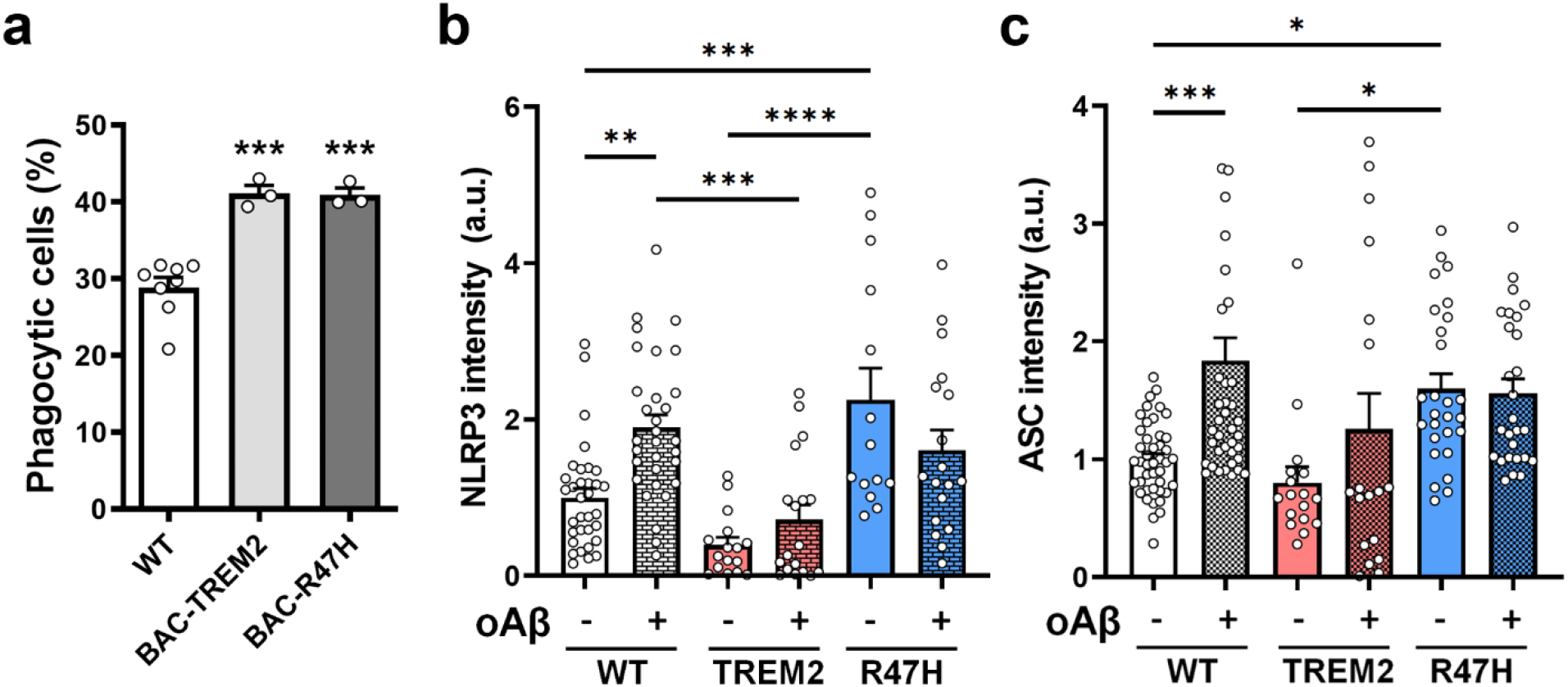
Overexpressing TREM2-R47H enhanced baseline phagocytic activity and attenuated oAβ-induced inflammasome activation in primary microglia. (**a**) Phagocytosis of Alexa488-conjugated microsphere by primary microglia were measured by flow cytometry. Phagocytic microglia were detected with strong fluorescent signal in the cells. (**b- c**) Primary microglia derived from WT, BAC-TREM2 and BAC-TREM2-R47H mice were treated with or without oAβ (5 μM, 24 h). Immunofluorescent imaging of NLRP3 (**b**) and ASC (**c**) were quantified. Average fluorescent intensity per cell was normalized to the non-treated controls of WT microglia. The graph is pooled results from 3 independent experiments. *n* = 3-4 per genotypes in each experiment. Data are represented as mean ± SEM. \**p* < 0.05, \*\**p* < 0.01, \*\*\**p* < 0.001, \*\*\*\**p* < 0.0001 compared to WT or otherwise indicated.

The NOD-, LRR-, and pyrin domain-containing 3 (NLRP3) inflammasome has an important role in microglial activation [52] and has been implicated in AD pathogenesis [53, 54]. NLRP3 in microglia can interact with the apoptosis-associated speck-like protein (ASC) to form an inflammasome complex known as ASC specks, which in turn cross seed and exacerbate amyloidosis [55]. Since loss of TREM2 enhances LPS-induced NLRP3 inflammasome signal [56], we next investigate whether elevating TREM2 or TREM2-R47H could differentially modulate NLRP3/ASC signaling in cultured primary microglia both at the baseline or upon challenge with oligomeric Aβ (oAβ; Fig. 3b, 3c). Interestingly, at the base line, we observed a significant elevation of both NLRP3 and ASC immunostaining in BAC-TREM2-R47H microglia compared to WT or BAC-TREM2 microglia, while the latter two genotypes did not show significant differences in the levels of these proteins. Upon oAβ administration, only WT microglia showed a significant increase in expression of NLRP3 and ASC compared to the baseline condition. BAC-TREM2 and BAC-TREM2-R47H microglia did not show a significant induction of these proteins from their respective baselines. The results suggest that overexpressing TREM2-R47H in microglia may elevate inflammasome activation at the baseline, while increased wildtype TREM2 or TREM2- R47H suppress inflammasome activation upon oAβ exposure. The latter is consistent with a shared anti-inflammatory effect of microglia overexpressing wildtype and R47H variant of TREM2 in response to oAβ, at least in an *in vitro* study.

Together, our *in vitro* studies revealed microglia overexpressing human TREM2-R47H retain the TREM2 function in enhancing phagocytosis and suppressing oAβ-induced inflammasome activation. Moreover, it also uncovers an intriguing effect of overexpressing TREM2-R47H, but not wildtype TREM2, in enhancing baseline inflammasome activation. However, this effect appears to be an *in vitro* phenomenon as evidence of microglial activation in BAC-TREM2-R47H were lacking in our *in vivo* studies (see below).

### Overexpressing TREM2-R47H does not change microglial morphology and interaction with amyloid plaques

In response to amyloid deposition, microglia migrate to and encircle the plaques and undergo morphological changes (termed microgliosis) including shortened and thickened processes and amoeboid shaped cell bodies [57]. We previously reported that overexpressing TREM2 in 5xFAD altered the morphology of plaque-associated microglia, which showed longer and more ramified processes while remaining encompassing the plaques. This phenotype is distinct from non-plaque-associated microglia or *Trem2*-deficient microglia. Here, we observed similar morphological changes of plaque-associated microglia in 5xFAD/TREM2 mice (Fig. 4). However, overexpressing TREM2-R47H in 5xFAD mice did not significantly alter the morphological responses of microglia in the presence of amyloid plaques (Fig. 4a, 4b, 4e-4g). Moreover, we observed similar number of microglia associated with the plaques in 5xFAD and 5xFAD/R47H mice, while the number of microglia associated with the plaques was reduced in 5xFAD/TREM2 mice (Fig. 4d). Together, our human BAC transgenic models reveal that TREM2-R47H exhibits a loss-of-function phenotype compared to wildtype TREM2 in altering microglial morphology and interaction with the amyloid plaques *in vivo*.

**Figure 4.**
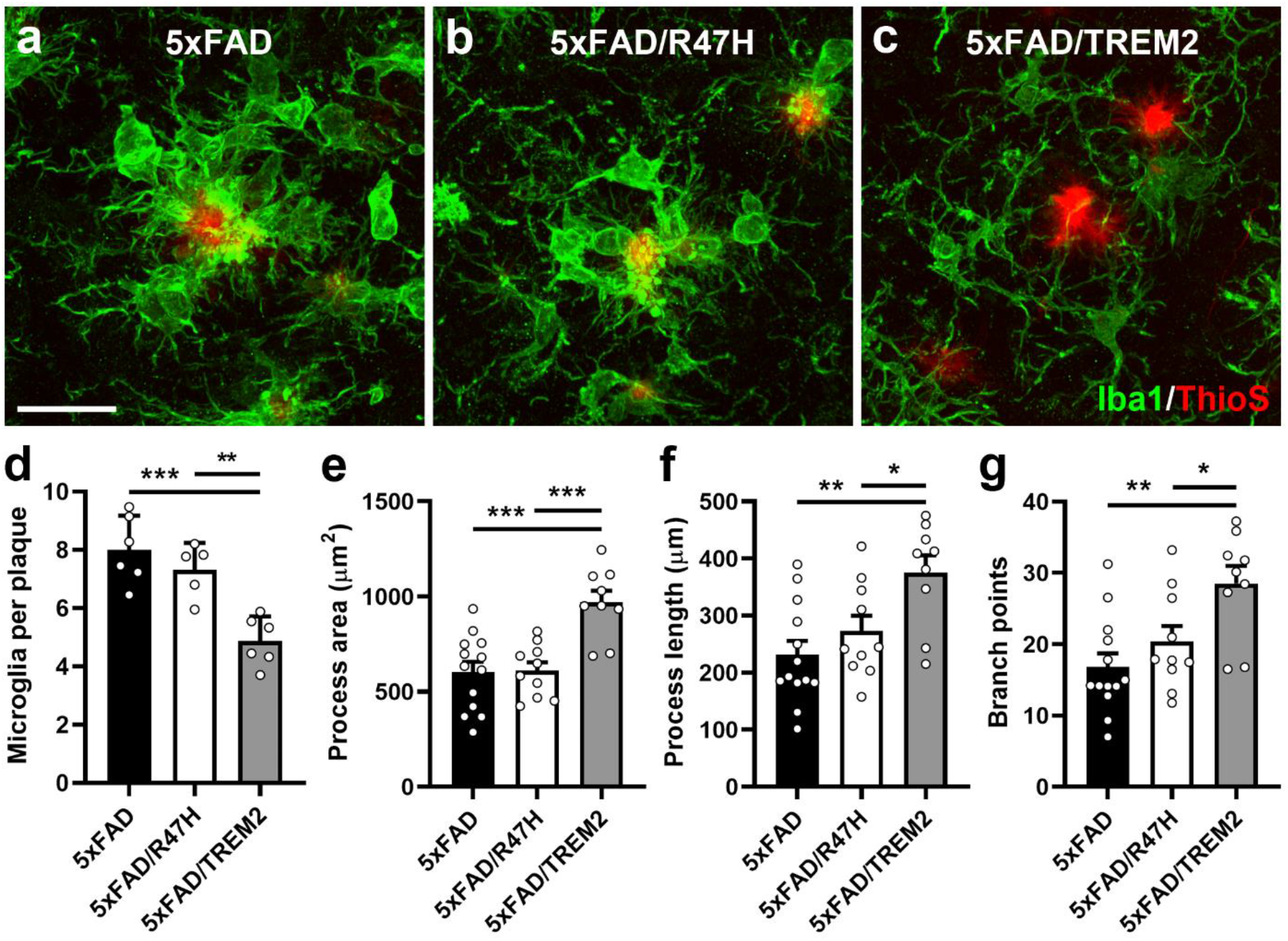
Overexpressing TREM2-R47H did not change the interaction of plaque-associated microglia and their morphology. **(a-c)** Representative images demonstrated the interaction between microglia (Iba1^+^, green) and the plaque (ThioS^+^, red) in 7 months old male 5xFAD (a), 5xFAD/R47H (b) and 5xFAD/TREM2 mice (c). Bar = 20 μm. **(d)** Amyloid plaques in cortex were randomly selected and Z-stack confocal images were taken for counting plaque-associated microglia (Iba1^+^) per plaque. *n* = 6 per genotype. **(e-g)** The morphological properties of plaque-associated microglia were measured by Imaris using z-stack confocal images taken under a 40X objective lens. The results are presented as total process area (e), process length (f) and branch numbers (g) per microglia. Images of a total of 1305 microglia from 192 plaques were analyzed and presented as mean ± SEM. *n* = 5 per genotypes, \*\*\**p* < 0.001, \*\**p* < 0.01, \**p* < 0.05.

### Increased TREM2-R47H expression failed to rescue cognitive deficits in 5xFAD mice

Our study thus far revealed that BAC-mediated increase of gene dosage of *TREM2-R47H* in 5xFAD mice could significantly reduce plaque load and remodel plaque types but was unable to reprogram plaque-associated microglial morphology or gene expression. The former is only a partial loss-of-function compared to 5xFAD with increased wildtype human *TREM2* gene dosage (Fig. 1), while the latter appears to be a full loss-of-function. We next assess whether cognitive behavioral phenotype of the 5xFAD mice is modified in 5xFAD/BAC-TREM2-R47H compared to 5xFAD, WT and BAC-TREM2-R47H littermates. To this end, we performed the contextual fear-conditioning (CFC) test, a hippocampus-dependent memory test that is compromised in 5xFAD mice [58]. Unlike increasing wildtype TREM2 gene dosage that significantly ameliorated the CFC deficits in the 5xFAD background [22], increased TREM2-R47H gene dosage did not reduce the CFC deficit of 5xFAD mice (Fig. 5). Noteworthily, we did not detect any CFC deficits in TREM2-R47H mice, suggesting overexpression of TREM2-R47H alone does not elicit impairment in this assay. Together, we conclude that the TREM2-R47H abolish the beneficial effects of wildtype human TREM2 in rescuing the behavioral impairment in 5xFAD mice.

**Figure 5.**
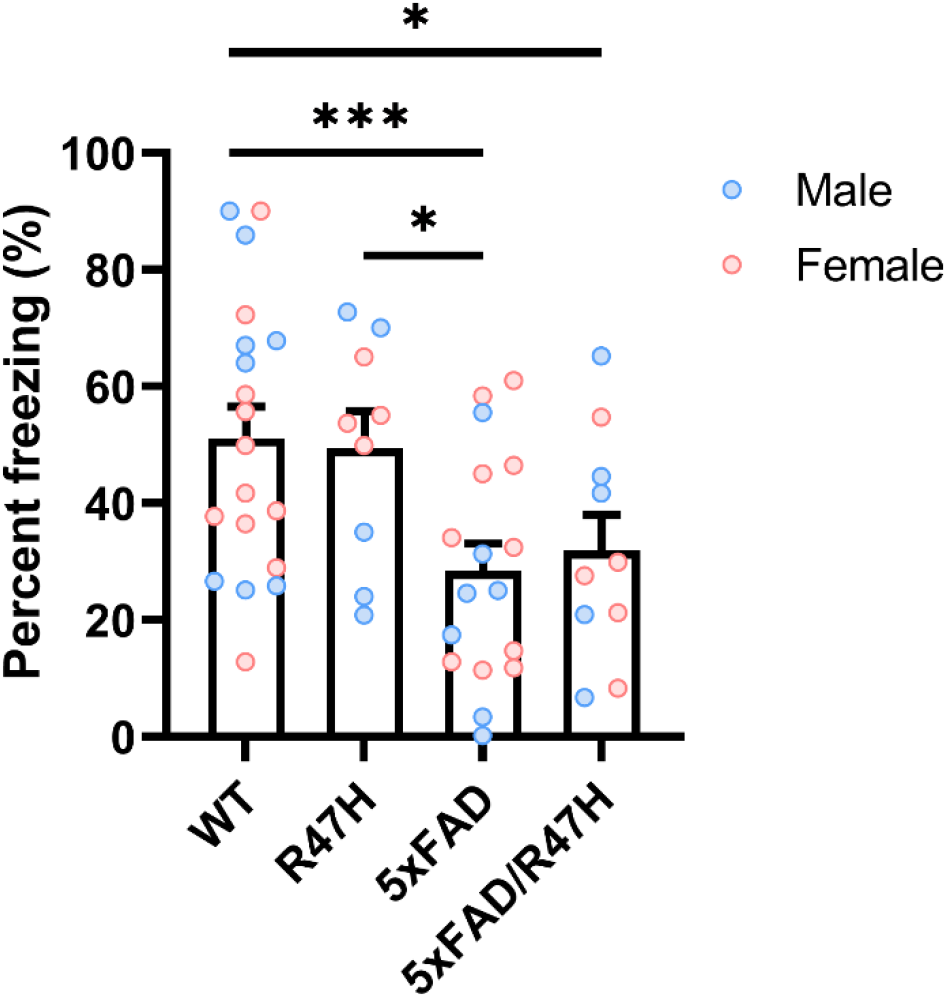
Increasing TREM2-R47H gene dosage failed to rescue cognitive deficits in 5xFAD mice. The contextual memory function of mice from the cohort of BAC-TREM2-R47H x 5xFAD (*n* = 10-19 per genotype) was evaluated by contextual fear conditioning and is presented as percent time freezing. Power analysis was performed to ensure >80% confidence levels with the number of animals used. \*\*\**p* < 0.001, \**p* < 0.05 compared to WT.

### Increased TREM2-R47H gene dosage did not modify cortical transcriptionopathy in 5xFAD mice

TREM2 plays an essential role in microglia recognition of Aβ pathology, including oAβ and amyloid plaques, and mediates a series of microglial responses, including barrier formation surrounding the plaque, morphological changes, and activation of disease-associated gene (DAM) expression signatures [25, 59, 60]. We previously demonstrated that overexpressing wildtype TREM2 reprogrammed microglial responsivity in mouse models of AD [22]. To evaluate the impact of TREM2-R47H expression on the molecular pathogenesis in 5xFAD mice, we performed RNA-sequencing (RNA-seq) with the cortical samples from WT, BAC-TREM2-R47H, 5xFAD and 5xFAD/BAC-TREM2-R47H male mice at 7m of age. We first performed principal component (PC) analyses and observed a well-defined separation based on 5xFAD genotypes (Fig. S2a). However, no clear separation could be detected based on the BAC-TREM2-R47H genotype even after corrected for the 5xFAD genotype (Fig. S2b). We next analyzed the number of significantly differentially expressed (DE) genes in the cortex across all four genotypes at 7m (FDR < 0.1; Fig. 6a and Table S1). We first replicated the robust cortical transcriptomic difference between 5xFAD and WT at 7m with 1,450 significant DE genes [22]. Additionally, we observed a strong positive correlation (Pearson’s *r* = 0.68) of DE gene Z statistics between 5xFAD vs. WT across the two studies, indicating a strong concordance and replicability of the two transcriptomic studies (Fig. 6b, S2c). The BAC-TREM2-R47H in either WT or 5xFAD backgrounds elicited few significant DE genes in the cortex, with only three DE genes between BAC-TREM2-R47H and WT and 6 DE genes between 5xFAD/BAC-TREM2-R47H vs 5xFAD (Fig. 6a). The latter is much fewer than the 54 DE genes in the cortical samples between 5xFAD/BAC-TREM2 and 5xFAD at this age [22].

**Figure 6.**
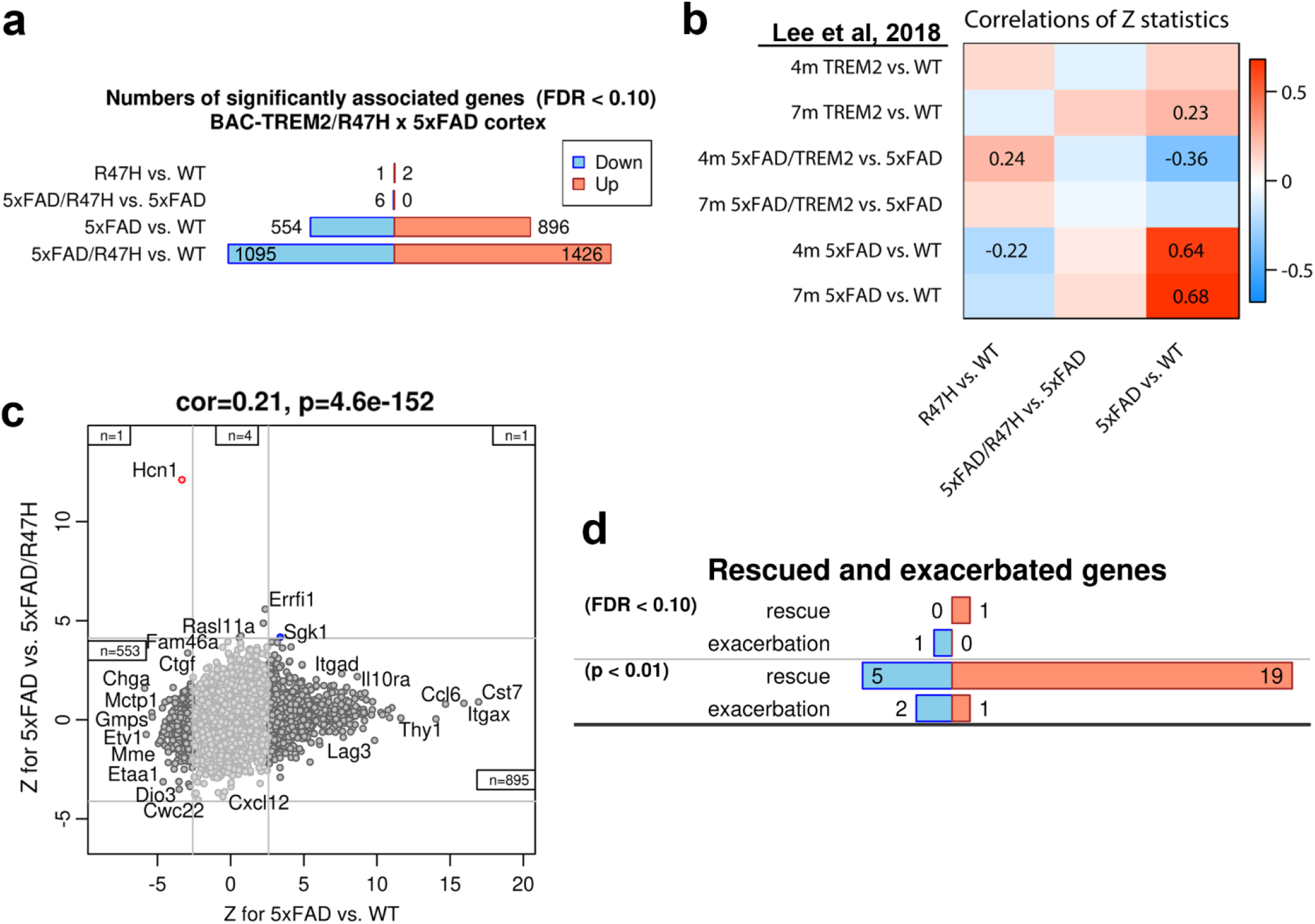
Differential gene expression analyses showed that overexpressing TREM2-R47H did not rescue transcriptomic changes in 5xFAD mice. (**a**) Numbers of significantly (FDR < 0.1) differentially expressed genes in the cortex of 7 months old male mice in genotype comparisons. Blue/red bars represent significantly down-/up-regulated genes. *n*=5 for WT, R47H and 5xFAD and n=8 for 5xFAD/R47H. (**b**) Heat map show the concordance of differential expression Z statistics of genotype pairs with the indicated genotype contrasts from our published BAC-TREM2 x 5xFAD dataset (Lee et al., 2018). Correlations whose absolute value is at least 0.2 are shown explicitly. (**c**) The transcriptome-wide “rescue plot” shows *Z* statistics for DE in 5xFAD vs. 5xFAD/R47H (y-axis) and 5xFAD vs. WT (x-axis) for all genes (each gene corresponds to one point). Rescued (concordant in this plot) and exacerbated (discordant) genes that pass the FDR threshold of 0.1 in both comparisons are shown in blue and red colors, respectively. Genome-wide correlations of Z statistics and the corresponding correlation p-values (for which the *n*=15,300 genes are considered independent) are indicated in the title of each panel. (**d**) Numbers of significantly rescued and exacerbated genes by BAC- TREM2-R47H at the two significance thresholds (FDR<0.1 and p<0.01). Blue/red bars represent genes that were down-/up-regulated for 5xFAD vs WT in 7 months old 5xFAD/R47H mice.

We next asked whether BAC-mediated expression of TREM2-R47H could rescue or exacerbate the transcriptome-wide dysregulation observed in 5xFAD mice. This transcriptome-wide “rescue/exacerbation” analysis was performed by correlating the Z statistics of 5xFAD vs WT in the X-axis and 5xFAD vs 5xFAD/BAC-TREM2-R47H in the Y-axis. A positive correlation indicates an overall transcriptomic rescue effect, while a negative correlation indicates an overall exacerbation [22]. We found the BAC-TREM2-R47H only elicit a weak rescue effect in 5xFAD mice (Pearson’s *r* = 0.21; Fig. 6c), which is much lower than that of BAC-TREM2 at 7m of age (Pearson’s *r* = 0.42, Lee et al., 2018). We next performed gene-level rescue exacerbation analysis (see Methods), and found BAC-TREM2-R47H did not significantly rescue or exacerbate any gene expression at FDR<0.1 (Fig. 6d). Even when we lower the threshold of significance to *p* < 0.01 or *p* < 0.05, no significant enrichment of those rescued and exacerbated genes was found (Table S2). Our previous study demonstrated that increasing wildtype *TREM2* gene-dosage in the 5xFAD mice elicited transcriptional reprogramming of microglial DAM gene expression, with downregulation of type 1 TREM2 gene-dose dependent (TD1) genes and upregulation of type 2 TREM2 gene-dose dependent (TD2) genes in 5xFAD/BAC-TREM2 compared to 5xFAD mice [22]. To compare the transcriptome-wide effects of BAC-TREM2-R47H or BAC-TREM2 in 5xFAD, we evaluated the association of Z statistics between 5xFAD/BAC-TREM2-R47H vs 5xFAD and 5xFAD/BAC-TREM2 vs 5xFAD, and we did not detect any significant correlation (Pearson’s *r* = -0.052; Fig. S3a). Consistently, expression levels of individual TD1 and TD2 genes, as measured by RNA-seq reads (Fig. S3b) and qRT-PCR (Fig. S4), were not significantly different between 5xFAD/BAC-TREM2-R47H and 5xFAD, nor between BAC-TREM2-R47H and WT. In summary, our cortical transcriptome study reveals that BAC-mediated expression of TREM2-R47H did not significantly alter the dysregulated transcriptome of 5xFAD, therefore showing a loss-of-function phenotype of TREM2-R47H compared to wildtype TREM2 in inducing the changes in microglial DAM gene signatures (i.e. downregulation of TD1 genes and upregulation of TD2 genes) in 5xFAD model [22].

### Selective modulation of TD1 and TD2 microglial gene transcripts in plaque-associated microglia by elevated expression of human TREM2 but not TREM2-R47H in 5xFAD model

Our previous findings on the opposing regulation of microglial TD1 and TD2 gene expression in 5xFAD/BAC-TREM2 compared to 5xFAD mice were based on the cortical bulk tissue RNA-seq [22]. However, it remains unclear how the coexpression and spatial distribution of TD1 and TD2 gene are in various microglial populations (e.g. homeostatic microglia or plaque-associated and activated microglia), and whether such distribution differs in BAC-TREM2- and BAC-TREM2-R47H-expressing microglia in 5xFAD models.

To quantitatively evaluate the spatial expression of TD1 and TD2 genes at cellular resolution and their relationships to the plaques, we performed multiplex *in situ* hybridization using the RNAscope (ACD Biosystems). We probed the brain sections from 7m old WT, 5xFAD, 5xFAD/BAC-TREM2 and 5xFAD/BAC-TREM2-R47H mice for the transcripts of human *TREM2 (hTREM2)*, two TD1 genes (*Tyrobp*, *Cst7*) and three TD2 genes (*Spp1*, *Gpnmb* and *Atp6v0d2*). Immunostaining was then performed following RNAscope to visualize the plaques (ThioS^+^ and 6E10) and microglia (Iba1^+^). We first confirmed the transcripts of *hTREM2* only present in 5xFAD mice carrying human BAC-TREM2 or BAC-TREM2-R47H transgenes, but not in 5xFAD or WT mice (Fig. 7, S5). We next quantified the percentage of plaque-associated microglia expressing the selected TD1/TD2 genes and their expression levels (see Methods). We found that a majority of plaque-associated microglia expressed *Tyrobp* and *hTREM2* in those 5xFAD mice carrying the BAC transgene. Indeed, *Tyrobp* and *hTREM2* were mostly co-expressed in the same microglia (Fig. 7a and Table S2). *Cst7* was also expressed in a majority of plaque-associated microglia in 5xFAD (86.4%) and 5xFAD/BAC-TREM2-R47H (72.8%), but only in 44.3% of plaque-associated microglia in 5xFAD/BAC-TREM2 mice (Fig. 6a, 6b). Importantly, the expression of expression of TD2 genes, i.e. *Spp1*, *Gpnmb* and *Atp6v0d2,* appears to be highly selective to the plaque-associated microglia in 5xFAD/BAC-TREM2 mice but not in the other two genotypes (Fig. 7a, 7b). Noteworthily, all the TD1/TD2 genes tested were exclusively expressed in the microglia associated with the plaques, but not in those away from the plaques in all three genotypes. Importantly, we observed heterogeneous TD1/TD2 gene expression patterns in plaque-associated microglia in all genotypes (Fig. 7a; Fig. S5b-S5c; Table S2-S3). For example, the level of *Cst7* expression varied even in the microglia of the same plaque in 5xFAD/BAC-TREM2 mice (Fig. 7a and 7b). These results highlight the potential heterogeneity in plaque-associated microglial responses at single-cell resolution, hence providing us a sensitive assay to study either *TREM2* or *TREM2-R47H* gene-dosage increase in modulating such response.

**Figure 7.**
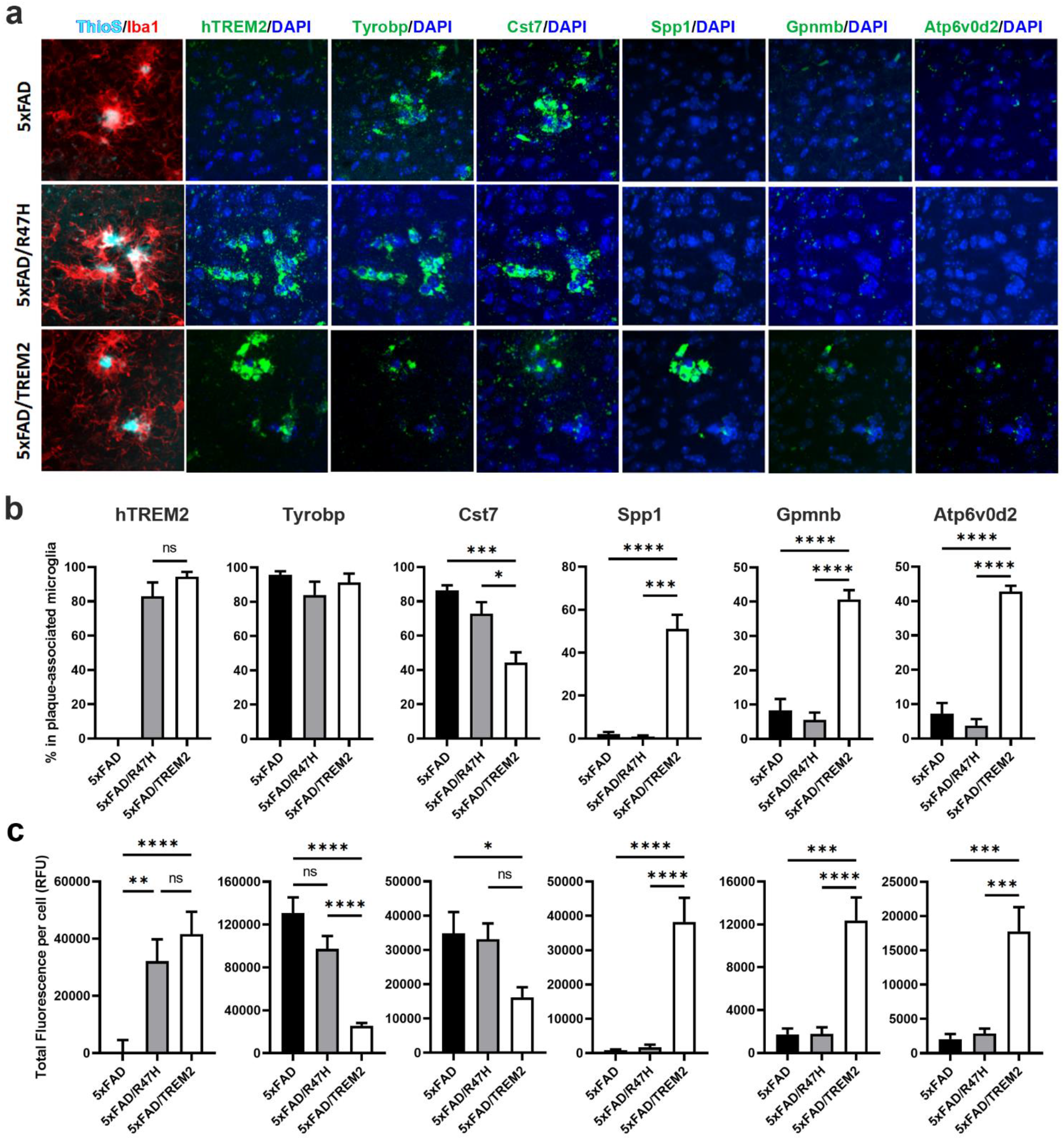
Overexpressing TREM2 and TREM2-R47H elicited distinct TREM2 dosage-dependent gene expression patterns in plaque-associated microglia. **(a)** HiPlex RNAscope was performed on matching cortical sections from 7-month-old male 5xFAD, 5xFAD/R47H and 5xFAD/ TREM2 mice for the expression of hTREM2, Tyrobp, Cst7, Spp1, Gpnmb and Atp6v0d2. The sections were then immunostained with anti-Iba1 to visualize microglia (red). Amyloid plaques and nuclei were stained with ThioS (cyan) and DAPI (blue), respectively. **(b-c)** The plaque-associated microglia expressing genes of interest were quantified and presented as percent positive cells over all plaque-associated microglia **(b)** and average total fluorescence per cell **(c)**. *n* = 5-6 per genotype. A total of 775 plaque-associated microglia was scored. Data are represented as mean ± SEM. \**p* < 0.05, \*\**p* < 0.01, \*\*\**p* < 0.001, \*\*\*\**p* < 0.0001

We next examined the relative levels of TD1 and TD2 genes in the plaque-associated microglia across the genotypes. We found that the TD1 genes (i.e. *Tyrobp* and *Cst7*) were highly expressed in plaque-associated microglia in 5xFAD and 5xFAD/BAC-TREM2-R47H mice, but significantly reduced in 5xFAD/BAC-TREM2 (Fig. 7a and 7c). In contrast, the transcripts for three TD2 genes (i.e. *Spp1*, *Gpnmb* and *Atp6v0d2*), which were expressed at low levels in plaque-associated microglia in the cortical sections of 5xFAD and 5xFAD/BAC-TREM2-R47H mice, were significantly upregulated in 5xFAD/BAC-TREM2 mice (Fig. 7a and 7c). No significant difference of microglial expression of TD1 and TD2 genes between 5xFAD and 5xFAD/BAC- TREM2-R47H mice was observed (Fig. 7c). Moreover, notwithstanding the possibility of probe sensitivity, it appears that only a smaller proportion of those *hTREM2+* microglia express *Gpnmb* and *Atp6v0d2* than those express *Spp1*.

We conclude that: (i). *TREM2* gene-dosage-dependent reprogramming of microglial gene expression, i.e. downregulation of TD1 genes and upregulation of TD2 genes, occurs selectively in the plaque-associated microglia; (ii). such reprogramming of plaque-associated microglial gene expression is abolished with the expression of *TREM2-R47H* variant, and hence the plaque-associated microglia of 5xFAD/BAC-TREM2-R47H has similar TD1/TD2 expression profiles as those in 5xFAD; and (iii). the upregulation of TD2 genes appear to be heterogeneous in the plaque-associated microglia in 5xFAD/BAC-TREM2 mice, a finding requires further validation.

## DISCUSSION

To study whether human TREM2-R47H could modify amyloid deposition and plaque-associated microglial response in an amyloid AD mouse model *in vivo*, we generated a novel BAC transgenic mouse model to increase human TREM2-R47H gene dosages. Unlike one of the murine knockin models of *Trem2-R47H*, which showed aberrant pre-mRNA splicing and nonsense-mediated mRNA decay [30, 31], the mRNA of *TREM2-R47H* in our new model was spliced properly and produced three isoforms as in human cells. Moreover, the expression of *TREM2- R47H* is at a comparable level as the wildtype *TREM2* in BAC-TREM2 mice, allowing a direct comparison to uncover the effects of the R47H mutation on TREM2 functions *in vivo*. In the 5xFAD background, our data suggested that the R47H allele appears to be a mixture of either partial or complete loss-of-function depending on the phenotypes examined, a finding that is consistent with the study of homozygous TREM2-R47H individuals who do not show PLOSL-like syndromes due to TREM2 deficiency [9]. Our study showed that increased TREM2-R47H levels significantly reduced the plaque load and shifted the types of plaques towards less filamentous, even though its effect size is significantly less than that of wildtype TREM2. In contrast, we found in 5xFAD model the increased expression of TREM2-R47H failed to recapitulate the effects of wildtype TREM2 in reprogramming the plaque-associated microglial morphological and transcriptomic phenotypes and overall amelioration of cognitive behavioral deficits. *In vitro*, we found that primary microglia overexpressing TREM2-R47H enhanced the baseline phagocytic activity to the level similar to overexpressing wildtype TREM2. Interestingly, the BAC-TREM2-R47H microglia had significantly higher NLRP3 and ASC levels than WT and BAC-TREM2 microglia. Yet, the levels of these inflammasome proteins were not significantly altered by oAβ stimuli, similar to BAC-TREM2 microglia. Overall, our study reveals that two functional aspects of human TREM2, i.e. plaque modification and Aβ phagocytosis vs morphological and transcriptomic reprogramming of plaque associated microglia, are differentially impacted by human TREM2-R47H mutation in the context of amyloid AD mouse brains.

A key strength of our study is from the design of the genetic construct. The BAC construct contains the entire human genomic locus of *TREM2*, allowing accurate, endogenous-like transgene expression [22]. The TREM2-R47H construct was modified from the TREM2 BAC, which was used to generate the BAC-TREM2 mouse model [22]. In the genome, there are several TREM-like genes with known function in innate immunity and possible AD modification located in the vicinity of TREM2 gene [10, 22, 61]. Thus, to interrogate and compare the function of human TREM2 and TREM2-R47H in mouse models of AD, we have deleted key exons of these other TREM-like genes so that only human *TREM2* or *TREM2-R47H* are expressed from the BAC transgenes [22]. To our knowledge, unlike other transgenic mouse models of human TREM2 or its R47H variant [32, 33], the BAC-TREM2 and BAC-TREM2-R47H models are the only models in which human TREM2 or TREM2-R47H are expressed under its genomic regulation and without expression of other human TREM-like genes. Our additional characterizations of our BAC models further reveal the advantages of these models to compare the differential functional roles of human TREM2 and TREM2-R47H in vivo: (i). both models have similar transgene copy number; (ii). both BAC-TREM2 and BAC-TRME2-R47H mice have the three human-like TREM2 alternative splicing variants; (iii). human *TREM2* and *TREM2-R47H* are selectively expressed in microglia in the brain; and (iv). the expression of *TREM2* and *TREM2-R47H* is selectively upregulated in plaque-associated microglia. Therefore, our current study provides a powerful novel human genomic transgenic mouse model system to study the AD-associated human TREM2-R47H variant *in vivo* to gain insights in AD pathogenesis and therapeutics.

Our study shed light on a functional role of TREM2-R47H variant in remodeling and reducing amyloid plaques *in vivo*. Prior studies revealed that *Trem2*-defficient microglia failed to encompass amyloid plaques and ultimately led to increased amyloid deposition and neurotoxicity [15, 21, 29]. In contrast, increasing *TREM2* gene dosage by a BAC transgene [22] as well as boosting TREM2 signaling with therapeutic antibodies [62] resulted in reduced amyloid burden and dystrophic neurites and ameliorated cognitive deficits in mouse models of AD. However, the impact of the TREM2-R47H variant on the modification of amyloid plaque phenotypes remains inconclusive. In one study, the “humanized” TREM2 and TREM2-R47H mice, in which BAC transgenes expressing either the common or R47H variant of human *TREM2* (and with both TREML1 and TREML2 genes on the BAC), were bred onto a murine *Trem2*-null background and then crossed with 5xFAD mice [33]. The study used 5xFAD/Trem2^-/-^ mice as a control and showed that *TREM2* common variant, but not *TREM2-R47H* variant, restored the microglial barrier functions around the plaque and activated microglial transcriptional signatures. However, this study did not address the plaque load or types in the brains of these mice, or any possible confounding effects with expression of other TREM-like genes from the BAC mice. In a recent study of novel murine *Trem2-R47H* knockin mice found that Trem2-R47H modulated the plaque load in 5xFAD at 4m in a sex-dependent manner; reducing ThioS+ plaques in male 5xFAD mice while increasing them in female mice [34]. The study also found reduced plaque intensity in the 5xFAD/Trem2-R47H mice compared to 5xFAD mice. These results suggest that Trem2-R47H may have a minor role in modifying plaque load or compaction. In light of these findings, our study represents strong in vivo evidence to date that the R47H variant of human TREM2 still retains the partial function of wildtype TREM2 in reducing plaque load in an amyloid AD mouse model *in vivo*.

Our in vitro study probed the mechanisms altered in microglia with elevated TREM2 or TREM2-R47H in response to Aβ. Prior studies showed that TREM2 could bind to and mediate phagocytosis of oligomeric Aβ [13, 15, 25, 63, 64]. We tested the idea of TREM2-R47H overexpression can boost in vitro microglia phagocytosis, similar to our prior findings with common variant TREM2 [22]. Indeed, we found that elevated expression of human TREM2-R47H significantly enhanced the baseline phagocytic activity in primary microglia compared to control microglia, and the level of increase is comparable to those microglia overexpression wildtype TREM2. Our in vitro finding is also consistent with the result that overexpression of both wildtype (or common variant) and R47H TREM2 restores defective phagocytosis of aggregated Aβ in Trem2-KO THP1 cells, suggesting the R47H allele may still retains TREM2 function related to phagocytosis [65]. Our study of inflammasome activation markers showed that wildtype microglia consistently showed oligomeric Aβ induced inflammasome activation (i.e. elevated NLPR3 and ASC expression), which is not found with TREM2 or TREM2-R47H overexpression. However, TREM2-R47H overexpression alone elicit the upregulation of NLPR3 even without oAβ challenge. Since oAβ inducted NLPR3 was found to drive pathogenesis in AD mouse models, including plaque deposition [53], our *in vitro* finding also suggests possible lack of Aβ-induced NLPR3 activation could support the plaque-reduction roles of *TREM2* and *TREM2-R47H* gene dosage increase, a hypothesis requires further investigation.

One important novel finding of this study is to show the effects of *TREM2* gene-dosage dependent reprogramming of plaque-associated microglial morphological changes and gene expression are completely lost with TREM2-R47H mutation. This finding adds further evidence to the interpretation that a critical role of TREM2 in the context of amyloid AD is to mediate the plaque-associated microglial response, both morphologically and transcriptomically, to contain and limit the plaque toxicity to surrounding neurons [17, 29, 66]. In terms of molecular signatures associated with the activated microglial state in AD mouse models [17, 67], our prior study showed that increasing TREM2 gene dosage in 5xFAD mice split such “DAM gene signature” into two subgroups: TD1 genes are upregulated by 5xFAD but partially reversed in 5xFAD/BAC-TREM2, while TD2 genes are upregulated by 5xFAD and further upregulated in 5xFAD/BAC-TREM2. In this study, we made two important observations with regard to the nature of TD1 and TD2 genes. First, in 5xFAD/BAC-TREM2 mice, top TD1 gene (i.e. Cst7, Tyrobp) and TD2 genes (*Spp1, Gpnmb and Atp6v0d2*), are predominantly expressed in plaque associated microglia and appropriately down-regulated (i.e. TD1) or up-regulated (i.e. TD2) by BAC-TREM2 as predicted by the bulk RNA-seq study [22]. Second, 5xFAD/BAC-TREM2-R47H failed to alter the 5xFAD transcriptomes (observed in the bulk cortical tissues) or reprogram the expression of TD1 and TD2 top genes in the plaque-associated microglia (observed by RNAscope). Current knowledge suggests both the altered expression of key TD1 and TD2 genes could be beneficial in the context of AD. *Cst7* is a TD1 gene that encodes an endosomal/lysosomal cathepsin inhibitor [68], and its upregulation is found in mouse models of ALS and AD [18, 22, 69, 70]. Increased expression of *Cst7* leads to microglial dysfunction by blocking lysosomal trafficking and subsequently impairs Aβ clearance, while reduction of *Cst7* alleviates lysosomal distress and enhances phagocytic activity of microglia and mitigates deficits in AD mouse models [71]. Spp1 (a TD2 gene) is a secretary matricellular glycoprotein significantly upregulated during the inflammation associated with AD and other neurodegenerative conditions [18, 70, 72]. In response to inflammation, Spp1 is secreted by microglia and acts through autocrine and paracrine signaling to activate and recruit microglia, macrophages, and astrocytes [73, 74]. It drives the activation microglial state that attenuates secondary neurodegeneration after ischemic stroke [74, 75], suggesting a protective role in the disease state. Together, our study unambiguously identifies the downregulation of certain pathogenic TD1 gene and upregulation of potentially protective TD2 genes in plaque-associated microglia upon increase of *TREM2* levels, and such reprogramming effects are abolished in the *TREM2-R47H* variant. Thus, our study uncovers molecular players that could mediate the dichotomizing effects of human *TREM2* and *TREM2-R47H* in AD pathogenesis, which could also serve as molecular markers to assess candidate therapies aiming to boosting TREM2 function or mitigating TREM2-R47H defects in AD.

## CONCLUSION

Our findings support that the AD-risk R47H allele of TREM2 is a partial loss of function mutation. Overexpressing TREM2-R47H by the BAC-TREM2-R47H transgene elicited partial yet significantly rescued amyloid pathology and enhanced microglial phagocytosis. However, TREM2-mediated transcriptional reprogramming was abolished. Our results also suggested that overexpressing wildtype TREM2 elicited a distinct subtype of plaque-associated microglia, which highly expressed TD2 genes and were not observed in 5xFAD and 5xFAD/BAC-TREM2-R47H mice. Overall, the study provided a mechanistic explanation for the pathogenic role of TREM2- R47H in AD and suggested that the novel BAC transgenic mouse models of TREM2 can be used for further mechanistic studies *in vivo* and translational research for AD therapy targeting TREM2.

## ABBREVIATIONS

AD: Alzheimer’s disease
ASC: apoptosis-associated speck-like protein
BAC: Bacterial Artificial Chromosome
BAC-TREM2: BAC-transgenic mouse model of TREM2
BAC-TREM2-R47H: BAC-transgenic mouse model of TREM2-R47H
CFC: contextual fear-conditioning
DAM: disease-associated microglial
DE: differential gene expression
GWAS: Genome-wide association studies
LOF: Loss-of-function
NHD: Nasu-Hakola disease
NLRP3: NOD-, LRR-, and pyrin domain-containing 3
NMD: nonsense mediated decay
oAβ: oligomeric Aβ
PC: principal component
PLOSL: polycystic lipomembranous osteodysplasia with sclerosing leukoencephalopathy
R47H: Arginine-to-Histamine mutation at amino acid residue 47
sTREM2: secreted TREM2
TD1: type 1 TREM2 gene-dose dependent gene
TD2: type 2 TREM2 gene-dose dependent gene
ThioS: Thioflavin S
TREM: Triggering Receptor Expressed in Myeloid Cell 2
12m: 12 months of age
4m: 4 months of age
7m: 7 months of age

## DECLARATIONS

## Acknowledgments

We acknowledge the help from UCLA Neuroscience Genomics Core (UNGC) for sequencing and reads alignment and UCLA Behavioral Testing Core (BTC) for advising on fear conditioning tests.

## Author Information

Center for Neurobehavioral Genetics, Semel Institute for Neuroscience & Human behavior, University of California, Los Angeles, Los Angeles, CA, USA; Department of Psychiatry and Biobehavioral Sciences, David Geffen School of Medicine, University of California, Los Angeles, Los Angeles, CA, USA; UCLA Brain Research Institute, University of California, Los Angeles, Los Angeles, CA, USA.

C.Y. Daniel Lee, Amberlene J. De La Rocha, Kellie Inouye, Peter Langfelder, Anthony Daggett, Xiaofeng Gu, Zoe Pamonag, Raymond G. Vaca, Jeffrey Richman, Riki Kawaguchi, Fuying Gao,

X. William Yang

Neuroscience anda Aging Research Center, Sanford-Burnham Prebys Medical Discovery Institute, La Jolla, California, USA

Lu-Lin Jiang, Huaxi Xu

## Author Contributions

CDL and XWY provided the conceptual framework, designed the experiments, and interpreted the results. CDL and XWY wrote the manuscript. AD generated the BAC-TREM2 and BAC-TREM2- R47H mice (Fig. 1a). XG contributed to the design of generating mouse models. CDL performed experiments and analyzed data shown in Fig. 1b-1d, Fig. 2, Fig. 3a, Fig. 4-8, and Supplemental Fig. S1-S5. KI performed experiment and analyzed data in Fig. 2, Fig. 4 and Supplemental Fig. S4. AJD, ZP and RGV contributed Fig. 6 and Supplemental Fig. S5. RNA-seq was performed by CDL and JR, and analyzed by PL, RK and FG. LJ and HX contributed to Fig. 3b and 3c. XWY provide funds for the study. All authors read and approved the final manuscript.

## Corresponding author

Correspondence to X. William Yang.

## Consent for publications

All authors have approved the contents of this manuscript and provided consent for publication.

## Availability of data and materials

All data generated in this study are included in this published article. Raw RNA-seq data is available through the gene expression omnibus (GSE number pending).

## Ethics approval and consent to participate

All our work was reviewed and approved by animal use committees at the University of California, Los Angeles.

