## Supplemental figures and tables for "BAC Transgenic Expression of Human TREM2-R47H Remodels Amyloid Plaques but Unable to Reprogram Plaque-associated Microglial Reactivity in 5xFAD Mice"

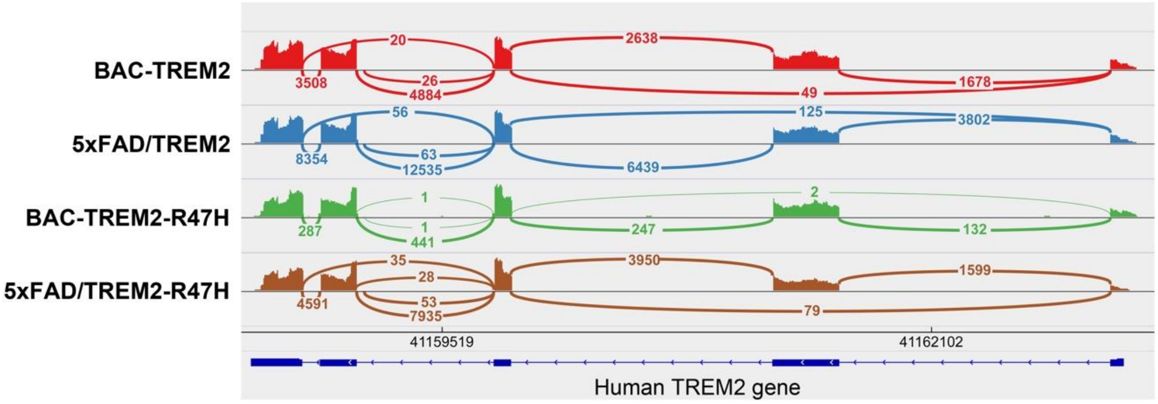


**Figure S1. Pre-mRNA of the TREM2-R47H transgene showed normal splicing pattern in both WT and 5xFAD background. Related to Figure 1.**
Splice variants with clearly observed alternative cassette-exon splicing events in the TREM2 locus of all 4 genotypes (n = 6). Curved lines indicated accumulative exon-exon junction spanning reads, with the number of associated reads centered on the line in the IGV Sashimi-Plot. The RefSeq mRNA exon transcript structure is shown below the plot.

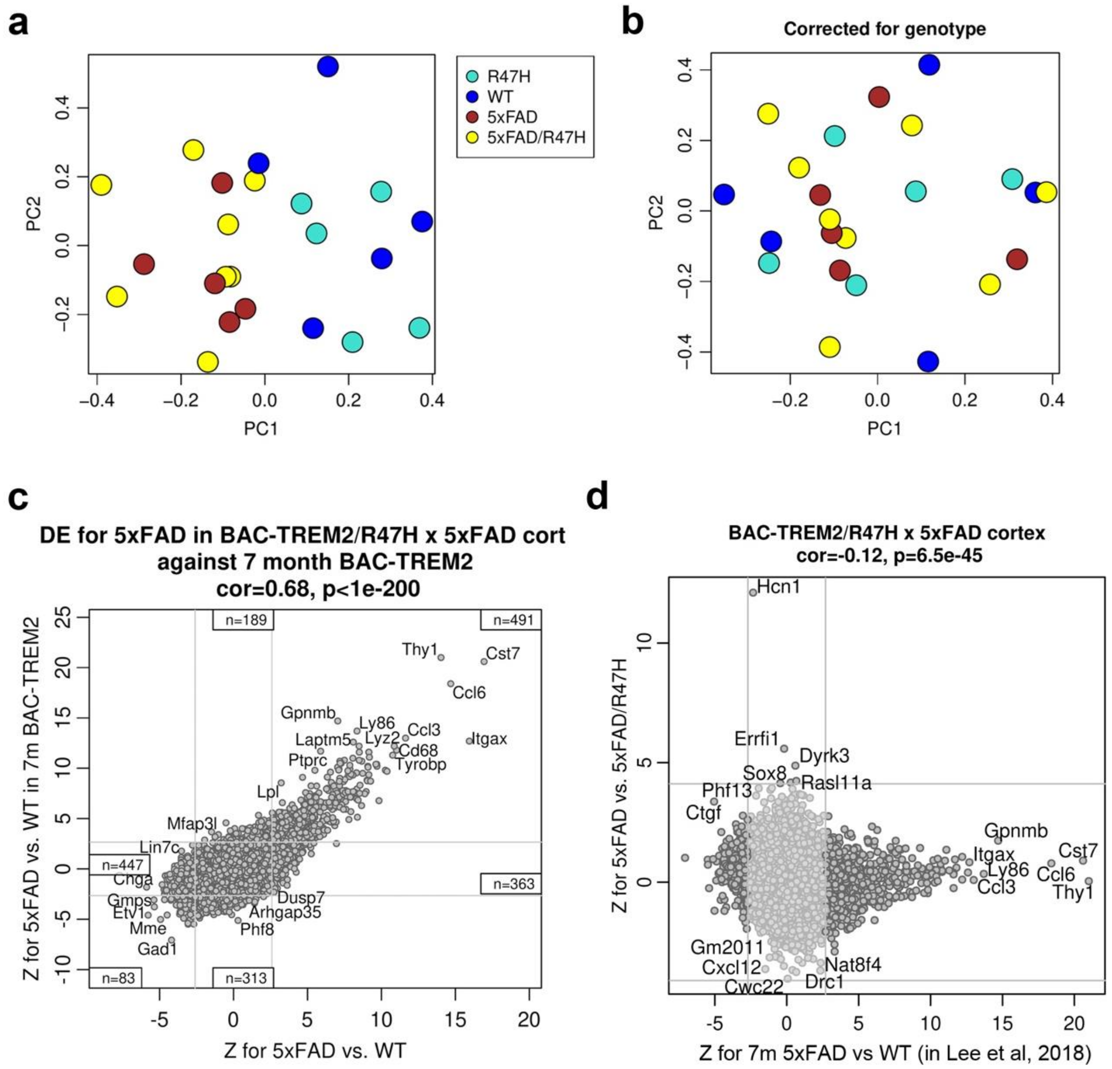


**Figure S2. Principle component analysis and concordance analysis of DE of the TREM2-R47H x 5xFAD cohort. Related to Figure 5.**

**(a-b)** The first two principal components of (**a**) the variance stabilized and outlier removed data (specifically, the scaled expression matrix for the top 8000 genes with highest mean expression) and (**b**) data from (a) corrected for genotype. Each dot represents the one mouse. The genotypes are indicated by color. **(c)** Z statistics of DE significance for 5xFAD vs. WT from 5xFAD/TREM2 (y-axis) and 5xFAD/R47H cohorts (x-axis). Each dot represents a gene. **(d)** *Z* statistics for DE significance in 5xFAD vs. 5xFAD/R47H (y-axis) from the 5xFAD/R47H cohort and 5xFAD vs. WT (x-axis) from the 5xFAD/TREM2 cohort (7 month-old; Lee et al, 2018) for all genes. Genome-wide correlations of Z statistics and the corresponding correlation p-values (for which the *n*=15,300 genes are considered independent) are indicated on the top of each plot.

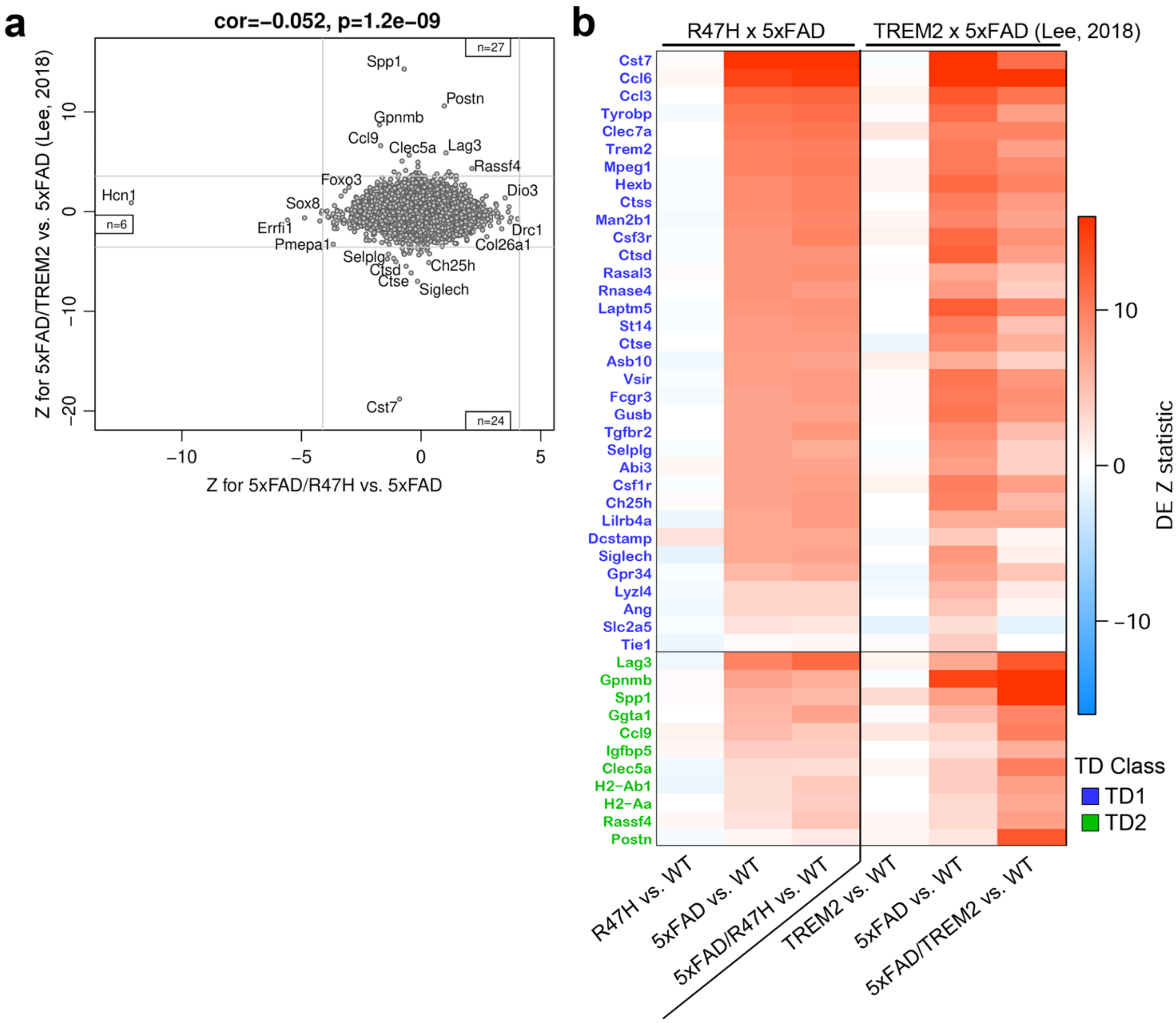


**Figure S3. The R47H mutation impaired TREM2-mediated transcriptional reprogramming of microglia without eliciting obvious gain-of-toxicity. Related to Figure 5.**
**(a)** DE significance *Z* statistics for 5xFAD/R47H vs 5xFAD and 5xFAD/TREM2 vs 5xFAD (dataset from Lee, 2018). **(b)** The heatmap shows DE significance *Z* statistics for selected TD1 (blue) and TD2 genes (green) in genotype contrasts listed below the heatmap.

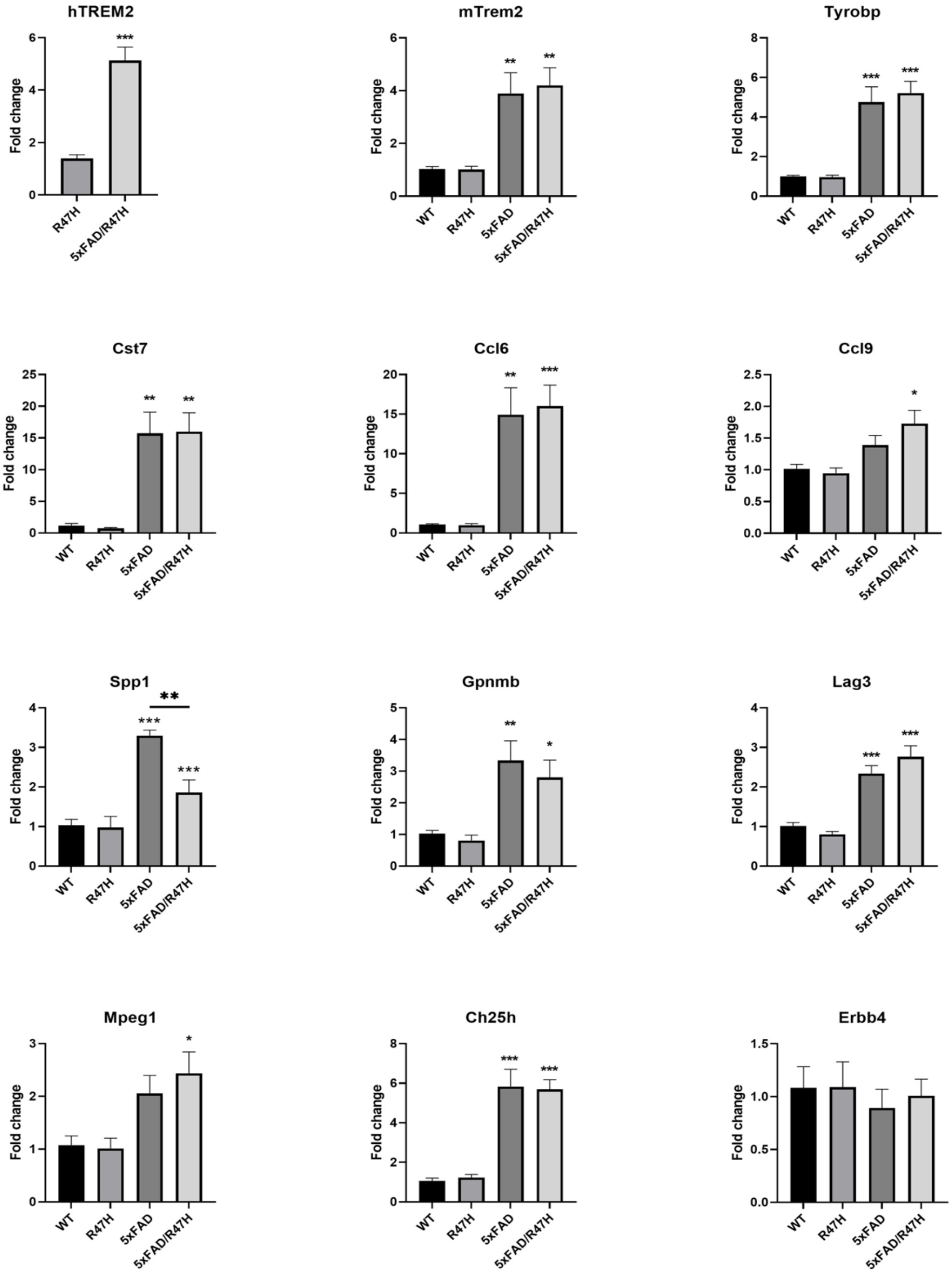


**Figure S4. Real-time PCR analysis to validate transcripts of selected TD genes. Related to Figure 5.**

Total RNA isolated from cortical tissues of 7 months old mice were reverse-transcribed to cDNA and followed by real-time PCR quantification with primers specific to human and mouse TREM2, and selected TD1 (Tyrobp, Cst7, Ccl6, Siglech, Ch25h and Mpeg1), TD2 (Spp1, Gpnmb, Lag3 and Ccl9) and TD3 (Erbb4) genes. Gapdh was used for loading control. The expression of transcripts was first normalized to internal Gapdh levels and then compared with WT. The levels of transcripts were presented as fold change over the WT controls. One-way ANOVA with Tukey post-hoc analysis was performed. The statistics are presented as comparing to WT controls unless specifically indicated in the graph. ***$p < 0.001$, **$p < 0.01$, *$p < 0.05$.

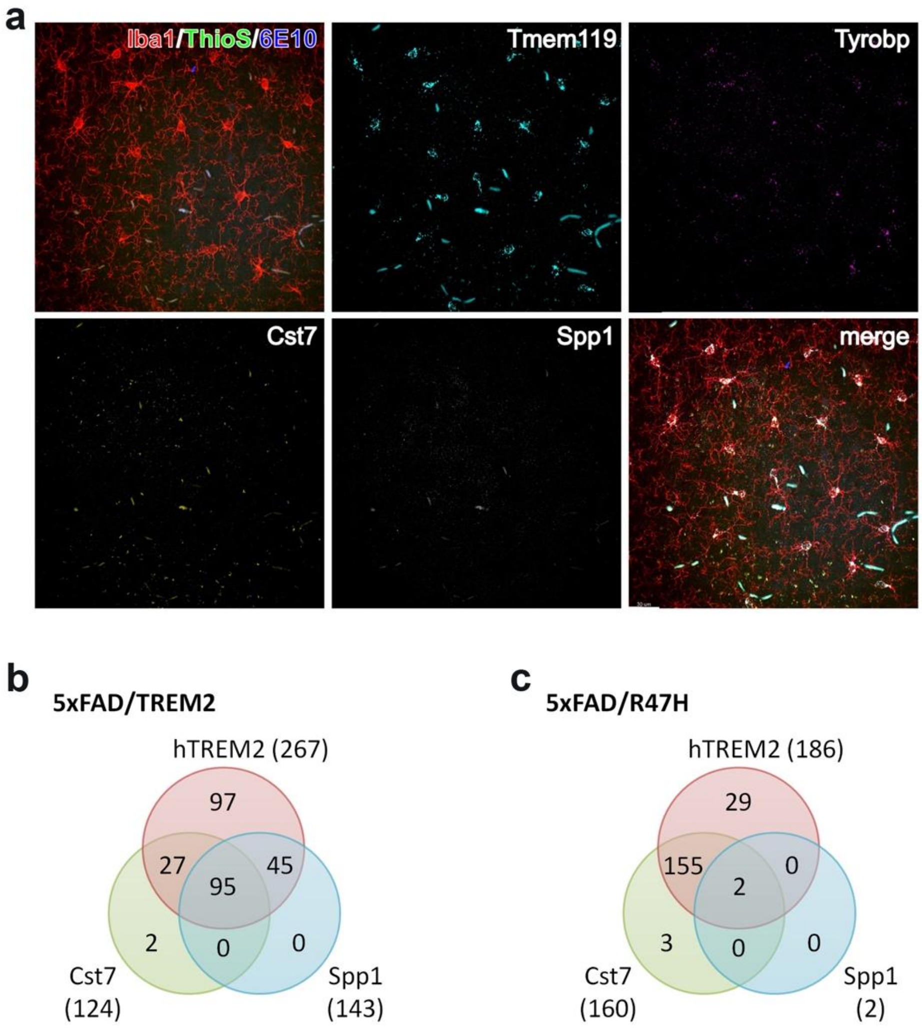


**Figure S5. RNAscope and gene co-expression analysis. Related to Figure 6.**
**(a)** Representative images of HiPlex RNAscope and subsequent immunohitochemistry performed on matching cortical sections (to Fig. 5a) from 7-month-old WT mice. Microglia were immunostained with anti-Iba1 (red). Amyloid plaques and nuclei were stained with ThioS (cyan) and DAPI (blue), respectively. Genes visualized by RNAscope are indicated at the upper right corners of the images. **(b-c)** Venn diagrams demonstrate the overlaps of hTREM2, Cst7 and Spp1 expression in the plaque-associated microglia of 5xFAD/TREM2 (b) and 5xFAD/R47H (c). Total numbers of cells expressing the specific genes are indicated in the parentheses.

| Category | Downregulated in 5xFAD vs. WT | Upregulated in 5xFAD vs. WT |
|---|---|---|
| **significant rescue ($p < 0.01$)** | **0** | **1** (Sall3) |
| **significant exacerbation ($p < 0.01$)** | **1** (Ctgf) | **4** (Nlrp3, Hs6st3, Ube2ql1, Mical1) |
| **significant rescue ($p < 0.05$)** | **0** | **14** (Sall3, Cebpd, Itgad, Npm3-ps1, Ldlr, Kcne1l, Il10ra, Ankrd13a, Cables1, Wdfy4, Aff1, Lfng, Msmo1, AI467606) |
| **significant exacerbation ($p < 0.05$)** | **19** (Ctgf, Arhgef3, Dusp14, Nat8f4, Hcn1, Drc1, Ly6g6e, Prss12, Brinp1, Plk2, Rai2, Rbm28, Cited2, Vmp1, Slit2, Pitpna, Abi1, Ptbp3, Amy1) | **30** (Nlrp3, Hs6st3, Cry2, Parp3, Rbfox3, Tia1, Fabp7, P3h3, Cry1, Rem2, Ube2ql1, Glt8d1, Phf13, Spata2, Wbp11, Sstr2, Fam196a, Mxi1, Thbs3, Hcn4, Crocc, Zfhx2os, Mical1, Dock6, Rassf4, Pom121, Mtmr11, Sec14l5, Itpkb, Uba7) |

**Table S2. Significantly rescued or exacerbated gene expression in 5xFAD/TREM2-R47H mice.**

The genome-wide correlations of DE Z statistics for 5xFAD vs. WT and 5xFAD vs. 5xFAD/R47H was used to define a measure of genome-wide rescue or exacerbation. The significantly rescued or exacerbated genes were also annotated as their differential expression pattern in 5xFAD vs WT. No significant enrichment was found in these groups of genes.

### 5xFAD (270 cells)

| | hTREM2 | Tyrobp | Cst7 | Spp1 | Gpnmb | Atp6v0d2 |
|---|---|---|---|---|---|---|
| **hTREM2** | **0** | - | - | - | - | - |
| **Tyrobp** | 0 | **255** | 90% | 2% | 6% | 5% |
| **Cst7** | 0 | 230 | **231** | 3% | 6% | 5% |
| **Spp1** | 0 | 6 | 7 | **7** | 43% | 29% |
| **Gpnmb** | 0 | 15 | 15 | 3 | **15** | 73% |
| **Atp6v0d2** | 0 | 12 | 12 | 2 | 11 | **12** |

### 5xFAD/R47H (224 cells)

| | hTREM2 | Tyrobp | Cst7 | Spp1 | Gpnmb | Atp6v0d2 |
|---|---|---|---|---|---|---|
| **hTREM2** | **186** | 97% | 84% | 1% | 5% | 4% |
| **Tyrobp** | 181 | **188** | 85% | 1% | 6% | 4% |
| **Cst7** | 157 | 159 | **160** | 1% | 7% | 4% |
| **Spp1** | 2 | 2 | 2 | **2** | 50% | 0% |
| **Gpnmb** | 10 | 12 | 11 | 1 | **12** | 33% |
| **Atp6v0d2** | 7 | 8 | 7 | 0 | 4 | **8** |

### 5xFAD/TREM2 (281 cells)

| | hTREM2 | Tyrobp | Cst7 | Spp1 | Gpnmb | Atp6v0d2 |
|---|---|---|---|---|---|---|
| **hTREM2** | **267** | 97% | 46% | 54% | 42% | 44% |
| **Tyrobp** | 259 | **261** | 47% | 54% | 43% | 45% |
| **Cst7** | 122 | 122 | **124** | 77% | 63% | 66% |
| **Spp1** | 140 | 138 | 94 | **143** | 65% | 65% |
| **Gpnmb** | 112 | 112 | 78 | 93 | **113** | 91% |
| **Atp6v0d2** | 117 | 117 | 82 | 93 | 103 | **119** |

**Table S3. Co-expression of TD genes.**
The expression of selected TD genes was visualized by RNAscope. Total cells analyzed in each genotype are indicated in the parentheses. The numbers of microglia co-expressed the gene pairs are labeled in the lower left half of the tables. Proportion of the co-expression is shown in the upper right half of the tables and color coded for better visualization. It is calculated as the percentage of cells co-expressing both genes in all cells expressing the gene indicated in the left column.
